# Individual differences in fear memory expression engage distinct functional brain networks

**DOI:** 10.1101/2025.05.12.653531

**Authors:** Barbara D. Fontana, Jacob Hudock, Neha Rajput, Dea Kanani, Dinh Luong, Justin W. Kenney

## Abstract

Fearful stimuli elicit a mix of active (e.g., evasion) and passive (e.g., freezing) behaviors in a wide range of species, including zebrafish (*Danio rerio*). Yet, how individuals vary in these responses, and how such differences are encoded in the brain, remains unclear. Here, we developed a contextual fear conditioning paradigm in which adult zebrafish learned to associate a specific tank with an aversive pheromone, conspecific alarm substance (CAS). Behavior during memory expression comprised of a mix of active and passive responses that clustered into four behavioral clusters: non-reactive, evaders, evading freezers, and freezers with the two high-freezing groups most clearly reflecting fear memory formation. We tested memory in four strains (AB, TU, TL, and WIK), and both sexes, finding that they shaped fear memory responses: males were more likely to increase evasion during CAS exposure, whereas TU fish were more likely non-reactive during recall. Whole-brain activity mapping during memory expression revealed that freezing engaged widespread neural activity that was particularly strong in the cerebellum, reticular formation, and parts of the telencephalon associated with olfactory processing. Evasion recruited visual threat detection regions and suppressed foraging and navigation circuits. Functional network analysis revealed that fish that froze, but had low levels of evasion (i.e., freezers) during memory expression had strong pallial-cerebellar connections and heightened interconnectivity between visual (pretectal) and thalamic like (preglomerular) nuclei. Animals that mixed freezing with evasive behaviors (i.e., evading freezers) had higher subpallial connectivity with autonomic and stress-regulatory regions like the hypothalamus and preoptic areas. Together, these findings demonstrate that distinct behavioral types during fear memory recall are supported by divergent brain-wide networks.

## Introduction

Fear is an emotional state elicited by perceived threat. Behaviorally, fear manifests as stereotypical responses that have a striking similarity across a range of taxa (Adolphs, 2013; Darwin, 1872). These responses can be classified into two broad categories: active (e.g., fighting or fleeing) and passive (e.g., freezing or immobility) with the decision of the right behavior at the right time being critical to survival. For example, evasive behaviors may be ideal when a predator is actively striking whereas freezing (i.e., a lack of movement) may be best when a predator is lurking. Recalling where fearful stimuli have been experienced is key to avoiding predation, enabling animals to anticipate danger and optimize survival strategies (Mobbs et al., 2015). Place-fear learning, termed contextual fear conditioning, has been widely studied, however, the extent to which individuals differ in fear expression and how such differences are represented in the brain, remains poorly understood. These questions are clinically relevant as variations in fear reactivity and fear memory are implicated in a variety of psychiatric disorders like post-traumatic stress disorder and anxiety (Mancini et al., 2025; Maren et al., 2013; Ressler et al., 2022).

One of the most reliable and ethologically relevant elicitors of fear in animals are alarm pheromones. These are small molecules released by animals as a form of communication, signaling the presence of a threat (Tirindelli et al., 2009). The existence of alarm pheromones is well established in many species of insects, fish, and mammals (Verheggen et al., 2010) and is increasingly recognized in humans (de Groot and Smeets, 2017). In fish, the first description of alarm pheromone was by Karl von Frisch (1938) in minnows (*Phoxinus laevis*), where he described how an extract of epithelial cells resulted in fear-related behaviors like freezing and erratic swimming. Responses were also species specific; fish preferentially respond to conspecific alarm substance (CAS) while ignoring substances from heterospecifics. He also found that fish formed memories of where they previously experienced exposure to CAS, and that individuals sometimes varied in their behavioral responses. More recently, variability in responsiveness to CAS has been reported in a range of fish species (Pfeiffer, 1977) including zebrafish (*Danio rerio*) (Speedie and Gerlai, 2008; Suboski et al., 1990). These findings suggest the presence of consistent individual behavioral differences in fear reactivity, but they have not been systematically examined.

Given the ethological relevance of CAS as a stimulus, it holds considerable potential for use in understanding individual variation in associative fear learning in zebrafish. In fear conditioning, a conditioned stimulus (CS) is paired with an intrinsically aversive unconditioned stimulus (US). Later presentation of the CS alone results in a conditioned response that is a measure of the strength of the memory. The CS can be either a discrete cue, like a light or sound, or a diffuse stimulus, like a tank or context. CAS has been successfully used as a US in Pavlovian fear learning assays in zebrafish where it has been associated with lights, odors, and contexts (Baker and Wong, 2019; Hall and Suboski, 1995; Maximino et al., 2018; Ruhl et al., 2017). These data suggest that fear learning using CAS is a robust phenomenon in zebrafish, making it ideal for determining if individual animals vary in fear memory expression and identifying the corresponding neural substrates.

To identify the presence of individual differences in fear behavior, we collected data from over 300 adult zebrafish across four inbred strains and both sexes. To increase throughput, we automated behavioral data analysis by combining DeepLabCut (Mathis et al., 2018) with a random forest supervised machine learning model (James et al., 2013). We then used unsupervised machine learning to identify the presence of individual differences in fear memory associated behaviors. Lastly, to identify the neural substrates of individual differences in fear memory, we mapped whole-brain activity of animals during fear memory recall using our recently developed pipeline for adult zebrafish (Rajput et al., 2025).

## Results

### Automated behavioral identification

To assess contextual fear memory in zebrafish we designed a three-day behavioral paradigm (**Figure 1A**). On the first day, animals were given five minutes to acclimate to the conditioning tank. On the second day, animals were placed back in the tank for 10 minutes. Baseline behavior was captured during the first five minutes (i.e., pre-exposure). CAS (derived from a mix of donors from all four strains and both sexes) or vehicle was then added to the tank for the last five minutes. On the third day, fish were placed back in the same tank in the absence of CAS to assess fear memory. To automate the identification of different swimming behaviors, we trained a DeepLabCut (Mathis et al., 2018) network to track three points on the fish: the head, trunk, and tail (**Figure 1B**). We then used the output from this tracking to train a random forest model to identify five different behaviors: straight swimming, normal turns, freezing, burst swimming, and erratic movements. We used 34 parameters derived from the tracking to capture the movement of the animals over time (**Table S1-1**). These features were calculated using a sliding window of 45 frames (750 ms) to capture the temporal dynamics of movement.

**Figure 1.**
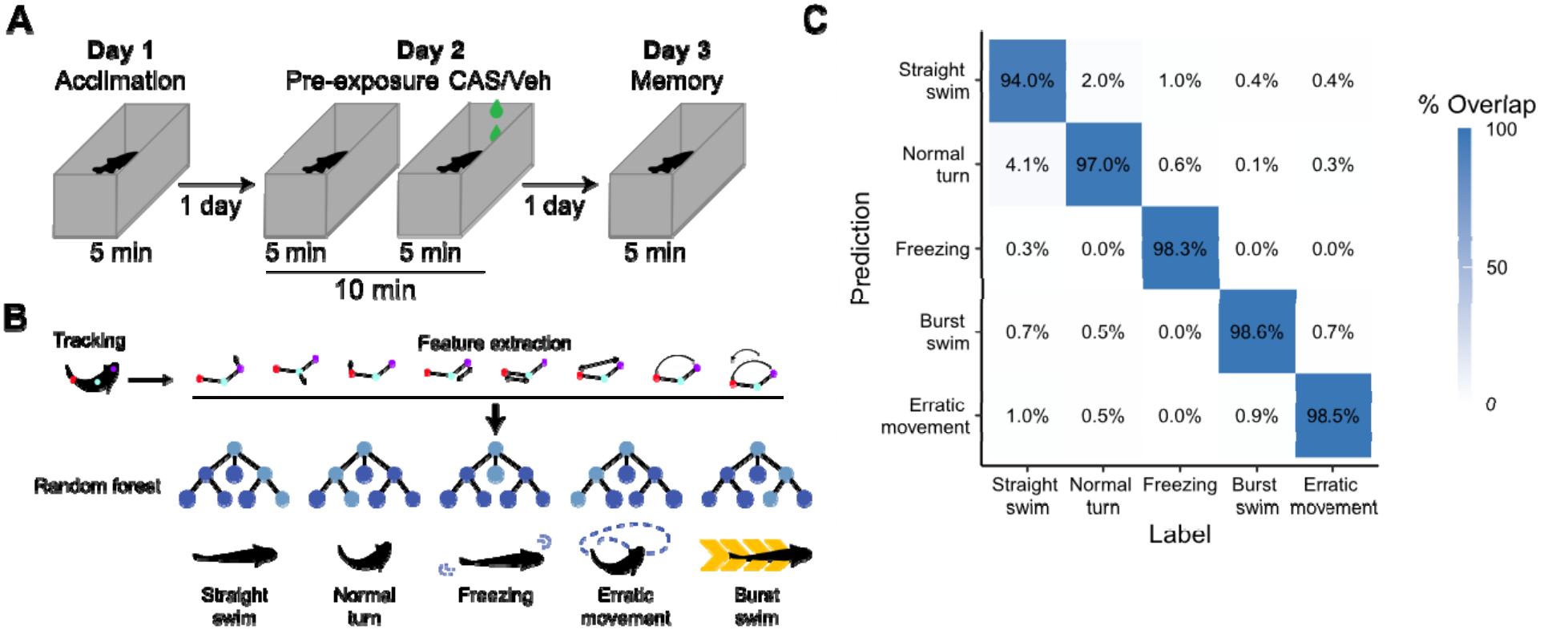
Fear memory paradigm and automated behavioral assessment. A) Adult zebrafish were trained in contextual fear conditioning over three days using CAS as the unconditioned stimulus. B) Automated behavioral assessment began with tracking using DeepLabCut, followed by feature extraction and then training of a random forest model for classification. C) Confusion matrix for the random forest model using a model trained on 80% of the data and tested on 20% of the data not used for training.

Approximately 14,000 frames were labelled for each of five different behaviors, with a 80/20 split for training/testing. We optimized the number of trees and number of features per tree, finding that 2,000 trees and 10 variables per tree minimized the out of bag error rate (2.88%). The resulting model had at least 94% accuracy for each behavior (**Figure 1C**). Notably, most of the misclassifications occurred when distinguishing between straight swimming versus normal turns and burst swimming versus erratic movements. Because these labels capture the same behavioral concept, and to simplify the subsequent analysis, we use “normal swimming” to refer to either straight swimming or normal turns and “evasive behavior” to capture erratic movement or burst swimming.

### Contextual fear learning

Given prior reports of behavioral variation in response to CAS, we hypothesized that this variation may be due, in part, to the background strain and/or sex of the animals. When looking at how fish respond to CAS, we found that almost all strains and sexes had an increase in freezing compared to vehicle treated animals (**Figure 2A**, top), and all groups were elevated relative to pre-exposure (**Figure 2B**, top). The only exception to this was female TL fish, where the vehicle treated animals also increased their freezing behavior, likely as a response to the small disturbance that occurs upon the addition of vehicle to the tank. We analyzed evasive behaviors as percent of time active (i.e., not freezing) since many fish spend a large proportion of their time freezing, giving us a better estimate of evasion that is less influenced by the more dominant freezing behavior. Evasive behaviors in response to CAS had more variation. Compared to vehicle treated animals, CAS treated males in two out of four strains increased evasive behavior (except WIKs and TUs that approached significance; **Figure 2A**, bottom), whereas there were no changes in females. Compared to pre-exposure, CAS resulted in an increase in evasive behaviors in all groups except male WIKs (**Figure 2B**, bottom).

**Figure 2.**
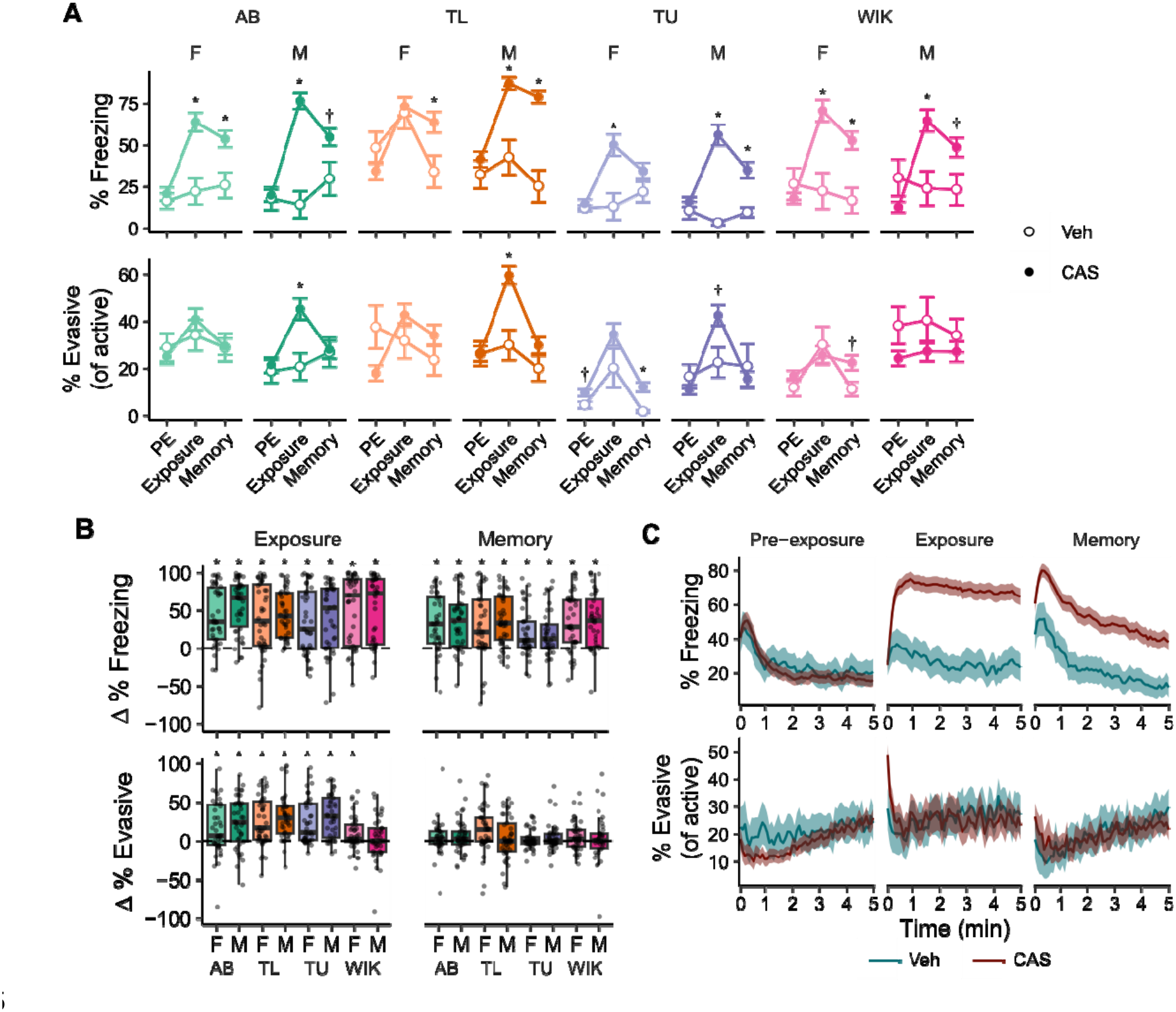
Behavioral responses during fear conditioning. A) Percent freezing (above) and % evasive behavior as a proportion of active behavior (below). Data are means ± SEMs. *-P ≤ 0.05, †-P ≤ 0.10 compared to vehicle treated fish via FDR corrected t-tests. B) Change (in comparison to pre-exposure) in percent freezing (above) and percent evasive behavior (below) during exposure to CAS (left) and memory day (right). *-P ≤ 0.05, compared to zero via FDR corrected one-sample t-tests. Boxplot center is the median, hinges are interquartile range, and whiskers are the hinge ± 1.5 times the interquartile range. C) Freezing (above) and evasive behavior as proportion of active behavior (below) over time during pre-exposure, exposure, and memory days. Lines are means and ribbons are 95% confidence intervals. Vehicle treated groups, n’s = 12; CAS treated groups, n’s = 40-42. Part C includes all the CAS-treated animals (N = 331).

We next asked whether fish formed a long-term associative memory between the tank and CAS. To test this, fish were placed back into the tank one day after CAS exposure with no CAS present (i.e., memory day). On memory day, there was elevated freezing compared to vehicle treated animals and relative to pre-exposure in nearly all strains and sexes, with the exception of female TU fish that did not differ from their vehicle treated counterparts and male WIKs and ABs that only approached significance (**Figures 2A** and **2B**, top). As before, evasive behaviors were more variable on memory day. When compared to vehicle treated animals, evasive behavior was only elevated in female TUs (**Figure 2A**, bottom). Compared to pre-exposure, no strains/sexes had an increase in evasive behaviors (**Figure 2B**, bottom).

To better understand the dynamics of freezing and evasive behavior, we examined temporal profiles of the behavior during the different stages of the paradigm (**Figure 2C**). During pre-exposure, there was an initial increase in freezing behavior that quickly decreased to ∼20%. Evasive behavior remained consistently low throughout the trial. During exposure, CAS-treated animals had an initial burst in evasive behavior that rapidly gave way to persistent freezing. On memory day, we saw an initial increase in freezing behavior in both vehicle and CAS-treated animals, but while the freezing in vehicle treated animals quickly declined, the freezing in CAS treated animals was higher and persisted for longer (**Figure 2C)**. Evasive behavior during memory day was largely similar between vehicle and CAS-treated animals, with only a slight increase in CAS treated animals at the beginning of the trial.

### Transitions between behavioral states

The aggregated data we have examined so far does not yield insight into how fish organize their behavior during the trials. To examine behavioral state changes, we constructed transition matrices by binning behavior into 750 ms bins that matched the sliding window used for behavioral identification (**Figure 3**). We found that at all stages, freezing is the most stable behavior (i.e., the behavior is most likely to follow itself), even during pre-exposure when it is a relatively small proportion of the overall behavior (**Figure 3A**). Amongst the normal swimming behaviors of straight swimming and normal turns, transitions are most likely between these states across all stages of behavior. A similar pattern is seen for evasive swimming (i.e., erratic movements and burst swimming); however, these behaviors are less likely to occur after normal swimming than normal swimming is to occur after evasive behavior. Overall, there are no large changes in the patterns of transitions across the different stages of the behavioral paradigm.

**Figure 3.**
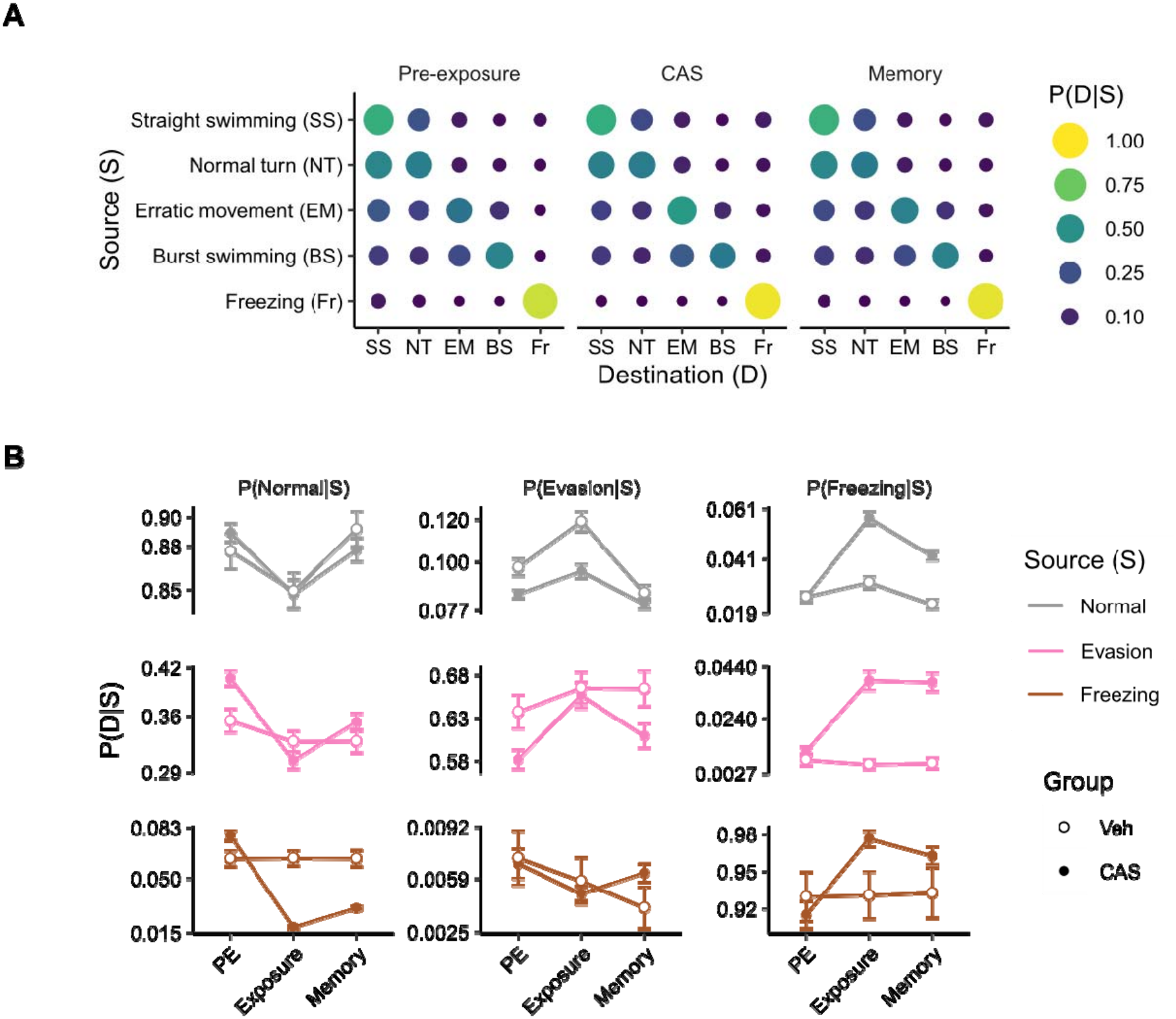
Transitions between behavioral states. A) Transition matrices capturing how often zebrafish change from the source to the destination state at different stages of the contextual fear memory behavioral paradigm. B) Transitions between source and destination states over time for normal, evasive, and freezing behavior. Error bars are 95% confidence intervals.

These data suggest that exposure to CAS does not result in wholesale reorganization of behavior, but instead a change in how much of each behavior is selected. To capture more subtle behavioral changes, we examined state transitions in response to CAS and at memory day in more detail (**Figure 3B**). To increase clarity, we collapsed straight and normal turns into “normal swimming” and burst swimming and erratic movements into “evasive behavior”. Here, we see that during both CAS and vehicle treatment, there is a slight destabilization of normal behavior such that normal behavioral bouts are shorter; however, in the vehicle group normal behavior is more likely to be followed by evasive behavior, and in CAS treated animals, normal behavior is more likely to be followed by freezing (**Figure 3B**, top row). This increase in evasion in the vehicle group may represent a response to the water disturbance that occurs when solution is added to the tank. We also see that CAS causes freezing behavior to occur more often and become more stable (**Figure 3B**, right column): compared to pre-exposure, freezing is more likely to occur following all behaviors at both exposure and memory day. Evasive behavior increases in likelihood following itself during CAS treatment, with little to no change following normal or freezing behavior (**Figure 3B**, middle column). This suggests that CAS results in longer bouts of evasive behavior without increasing its likelihood of occurring. In contrast, the CAS-induced elevation in freezing behavior during exposure and memory appears to be due to an increase in both the duration and likelihood of freezing bouts.

### Individual differences in fear memory expression

Given the large amount of variation in behavioral responses (**Figure 2B**), we hypothesized the presence of distinct fear behavioral types. We clustered the behavioral data using memory day because there was greater variation due to the lower levels of freezing compared to CAS exposure. To uncover behavioral types, we applied Louvain clustering to a series of k-nearest neighbor networks (KNNs) and identified the k-value that yielded robust cluster number (i.e., did not change over a wide range of k’s) that maximized internal clustering metrics (k = 74; **Figure S4-1A**) (Rajput et al., 2022). Distances between nodes (i.e., individual fish) in the network were calculated using percent evasion, percent evasion of active swimming, and percent freezing. Percent evasion of active swimming was included because fish that exhibit high levels of freezing will necessarily be low on non-freezing behaviors. By calculating evasion as a percent of active (i.e., non-freezing) behavior, we better capture what fish choose to do when not freezing. From this analysis, we identified four behavioral clusters (**Figure 4A**): (1) “non-reactive”: fish low in evasion and freezing; (2) “evaders”: fish low in freezing and high in overall levels of evasive behavior; (3) “evading freezers”: fish high in freezing and high in evasion as percent time active behaviors; (4) “freezers”: fish high in freezing and low in evasive behaviors (**Figure 4B**).

**Figure 4.**
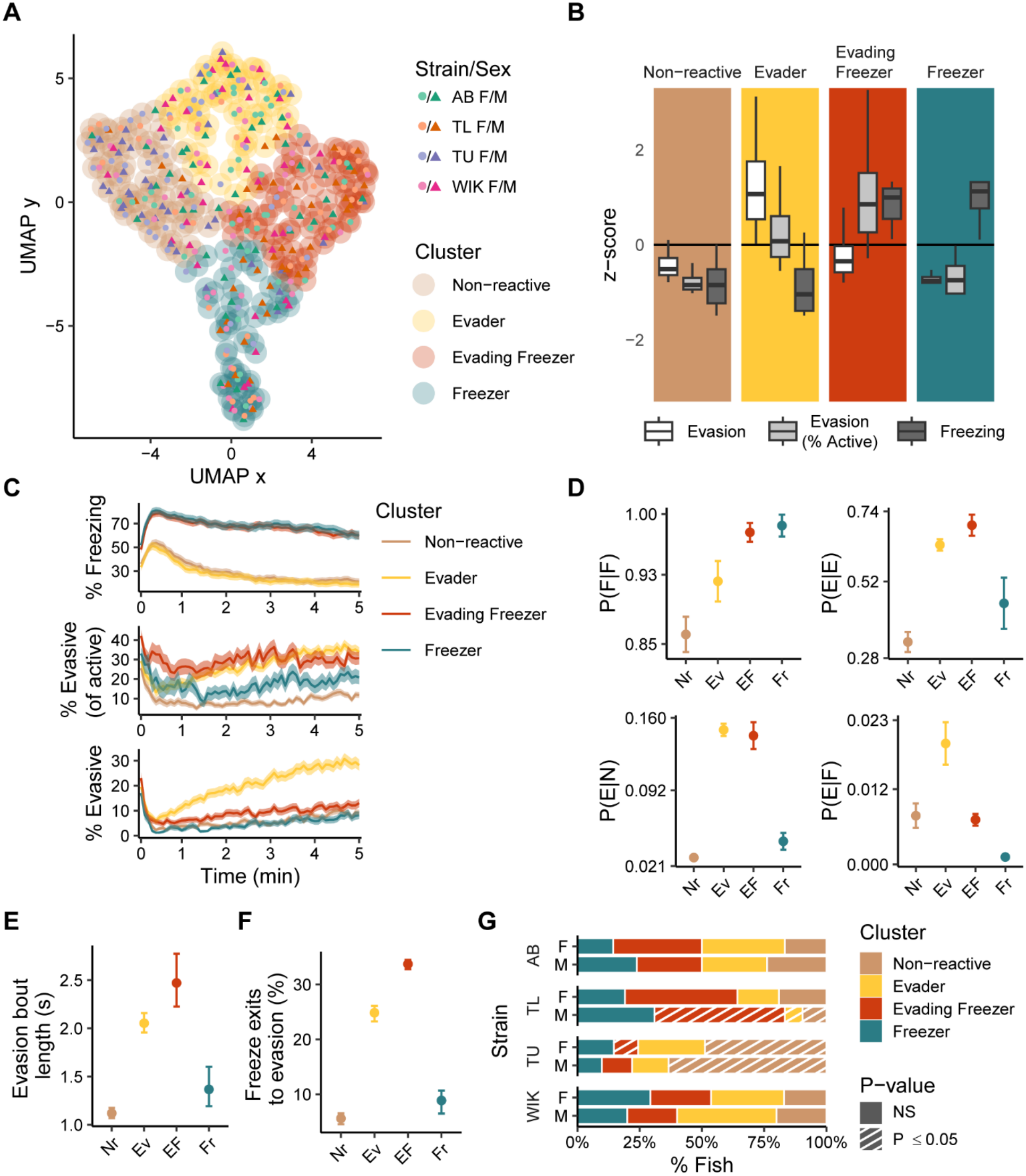
Clustering of zebrafish behavior during fear memory recall. A) Two-dimensional representation of the behavioral space used for clustering using a uniform manifold approximation (UMAP). The outer circles delineate the clusters as defined using Louvain clustering applied to a k-nearest neighbor network (k = 74). Each point represents an individual fish (N=331). B) Behaviors associated with the identified clusters presented as z-scores. Box plots indicate median (center line), interquartile range (box ends) and ± 1.5 times the interquartile range (whiskers). C) Behaviors across time for the different clusters; ribbons indicate 95% confidence intervals. D) Transition probabilities [P(Destination | Source)] for a subset of behaviors across the four identified clusters for a subset of transitions (full set of transitions can be found in Figure S4-3). F = freezing, E = evasion, N = normal behavior. Nr = non-reactive, Ev = evaders, EF = evading freezers, and Fr = freezers. Error bars are 95% confidence intervals. E) The average evasion bout length derived from P(E|E) transitions. Error bars represent 95% confidence intervals. F) Percentage of time fish exit freezing to evasion (instead of normal swimming) derived from P(E|F) and P(E|N). Error bars represent 95% confidence intervals. G) Percentage of animals falling into each behavioral cluster across strain and sex. Striped bars represent under/over representation using permutation tests with FDR corrections (P ≤ 0.05); n’s = 40-42 per strain and sex.

We also explored two other variable combinations for identifying behavioral clusters, finding them both to be suboptimal compared to the three variable approach above: (i) using percent evasion and percent freezing, and (ii) using isometric log ratios to generate a 2-D orthonormal basis from percent freezing, percent evasion, and percent normal swimming taking into consideration the compositional nature of our data (Aitchison, 1982; Egozcue et al., 2003; Greenacre, 2018). In both cases, the stable range of k’s in the KNN yielded four clusters. Two of these clusters mapped onto prior clusters (non-reactive and evaders) and two where freezers were split into medium and high groups (**Figure S4-2**). However, the robust k-regime for both of these alternative approaches did not overlap with maximal internal clustering metrics (**Figures S4-1B & C**), making them less optimal than our three-variable approach (**Figure S4-1A**).

Temporal profiles of behavior also support the presence of four distinct groups (**Figure 4B**). For example, although both freezer groups overlapped in the percent freezing behavior over time, they differ in their evasive behavior, where the evading freezers have higher levels of active behavior spent evading throughout the entire memory trial (**Figure 4C, middle**). Although the overall levels of evasion in evading freezers are considerably lower than evaders, and only just above freezers, (**Figure 4C, bottom**), this is an artifact due to the high levels of freezing in this group that leaves less time for other behaviors.

Behavioral state transitions are an independent way to determine if the clusters we identified are distinct since these transitions were not incorporated into the machine learning. This analysis also supports the distinction between evading freezers and freezers (**Figure 4D-F** and **Figure S4-3**). For example, although freezers and evading freezers overlap in freezing bout length [P(F|F)], they significantly differ in evasive bout length [P(E|E)] (**Figure 4D**) where the average bout length of evading freezers [2.77 s, 95% CI (2.22, 2.47)] is 60% more than freezers [1.60 s, 95% CI (1.19, 1.37)] (**Figure 4E**). Furthermore, the likelihood that evading freezers transition from freezing to evasion [P(E|F)] is six times that of freezers (**Figure 4D**), resulting in evading freezers shifting from freezing to evasion 33% of the time versus 9% in freezers (**Figure 4F**). Thus, despite a small difference in overall evasion, freezers and evading freezers organize their behavior differently, consistent with being distinct behavioral types.

Finally, we asked whether fear memory behavioral types are affected by the strain and/or sex of the fish (**Figure 4G**). We found that in AB and WIK fish there was neither over nor underrepresentation in any of the four behavioral types. In contrast, TU fish were overrepresented in the non-reactive group, and TL males were overrepresented in the evading freezer group. This suggests that TU fish may be a poor choice for studying CAS-induced fear learning.

### Behavioral clusters over time

We next asked if behavior during pre-exposure or CAS exposure is predictive of behavior at memory day. To do this, we identified which cluster each fish fell into at each behavioral stage (**Figure 5A**). In the vehicle treated group, animals were largely identified as either non-reactive or evaders at all three time points. In CAS-treated fish, only 14% of animals belonged to one of the freezing groups at pre-exposure. During CAS exposure, this proportion increased to 66%. On memory day, 48% of CAS treated fish were in one of the two freezing groups, about equally split between evading freezers (28%) and freezers (20%). We found that there was a weak relationship between behavior during pre-exposure and memory day (**Figure 5B**). Evading freezers on memory day were just as likely to come from pre-exposure evaders (41%, 39/94) as from non-reactive fish (38%, 36/94), with nearly half (46%) arising from non-evading groups at pre-exposure (non-reactive or freezers). If we track forward transitions from pre-exposure to memory, we find that baseline non-reactive fish as likely to be evading freezers (23%, 36/157) as freezers (22%, 35/157). However, baseline evaders had a bias towards becoming evading freezers (30%, 39/129) on memory day instead of freezers (14%, 18/129). Thus, animals exhibiting the evading freezer phenotype reflect a combination of acquired responses and baseline traits.

**Figure 5.**
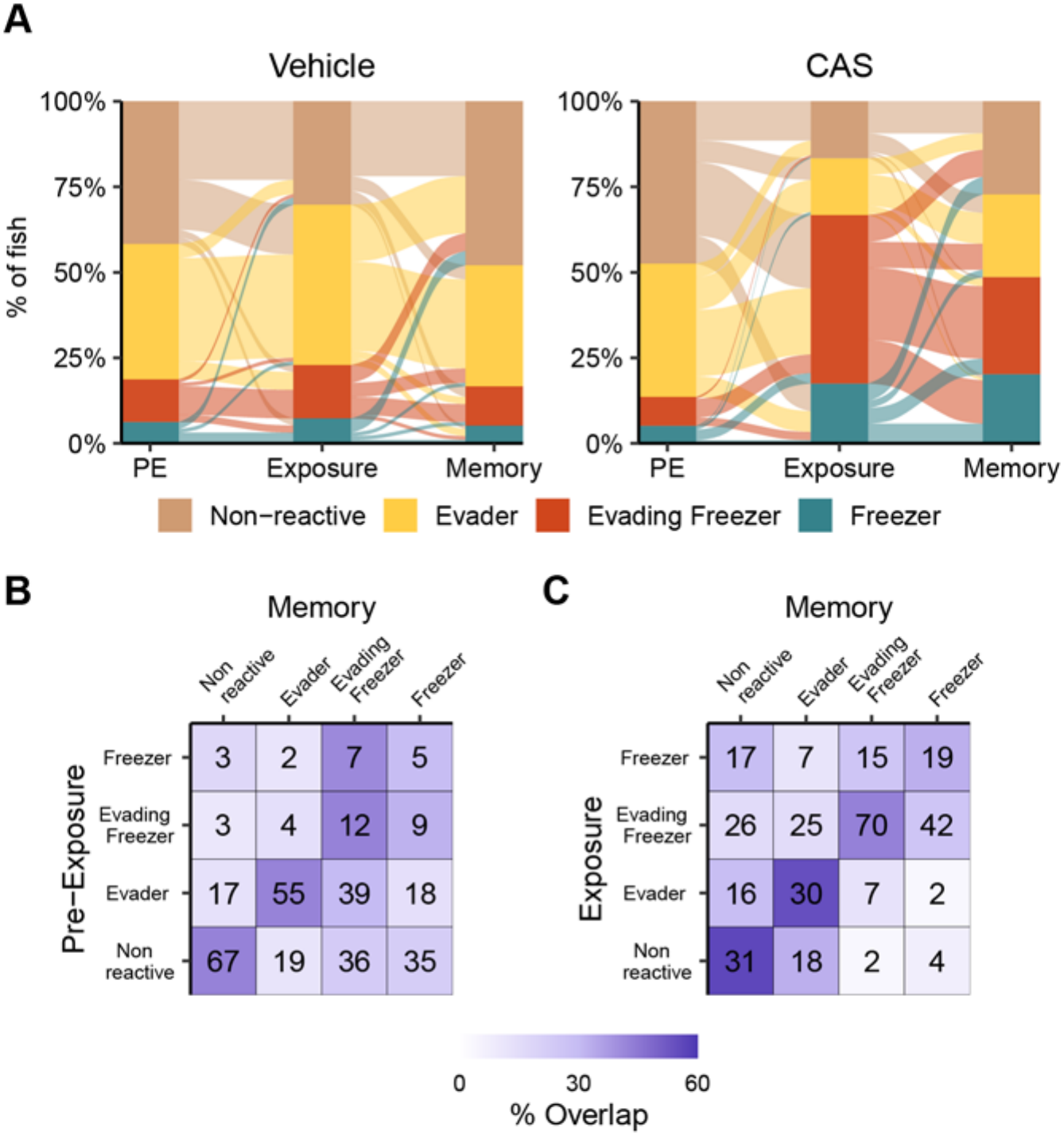
Transitions between behavioral clusters over time. A) Alluvial plot indicating the number of fish in each cluster during pre-exposure, exposure, and memory day for animals exposed to vehicle (left) or CAS (right). B) Confusion matrix of fish in different clusters during pre-exposure and memory day. C) Confusion matrix of fish in different clusters during exposure and memory day. Percent overlap is the proportion of fish at pre-exposure or exposure day that were in the same cluster at memory day. Vehicle treated fish; N = 96; CAS treated: N = 331.

We found that the behavioral cluster of the fish on exposure day was better at predicting behavior on memory day (**Figure 5C**, diagonal). Most fish in the non-reactive and evader groups on memory day fell into the same category during exposure (54-56%) or transitioned between evaders and the non-reactive groups (29%). Evaders and non-reactive fish rarely transition to freezing groups on memory day (11-16%). In contrast, a substantial minority of animals in the freezing groups transitioned to non-freezing groups (30-41%) with the majority remaining in one of the high freezing clusters (58-69%). Taken together, these data suggest that behavior at memory day is better predicted by behavior in response to CAS than prior to exposure to the stimulus, although there remains a sizable minority of fish that respond to CAS but do not appear to form a memory.

### Neural basis of freezing and evasion

To determine the brain regions underlying different behaviors, we performed whole-brain activity mapping followed by partial least squares (PLS) analysis. We chose the AB strain of fish because they showed approximately equal distribution of the different behavioral types (**Figure 4G**). For brain mapping, animals were euthanized 15 minutes after fear memory recall and processed for tissue clearing using iDISCO. Neural activity was captured using *in situ* hybridization chain reaction (HCR) against *cfos* mRNA. Brains were imaged using light-sheet microscopy and then registered to the adult zebrafish brain atlas (AZBA) (Kenney et al., 2021; Rajput et al., 2025). We used PLS analysis (Krishnan et al., 2011) to determine how brain activity covaries with behavior during fear memory recall (**Figure 6**). This approach uncovers latent vectors that capture the maximal covariance between neural and behavioral states. The first behavioral contrast identified was significant (P = 0.0090), capturing freezing in the positive direction and evasion/normal swimming in the negative (**Figure 6A**, left); the second contrast approached, but did not reach significance (P = 0.107; **Figure 6A**, right). However, because this second contrast uniquely captured evasive behavior, we analyzed it with the caveat that these findings are not as strongly supported as those for the first contrast.

**Figure 6.**
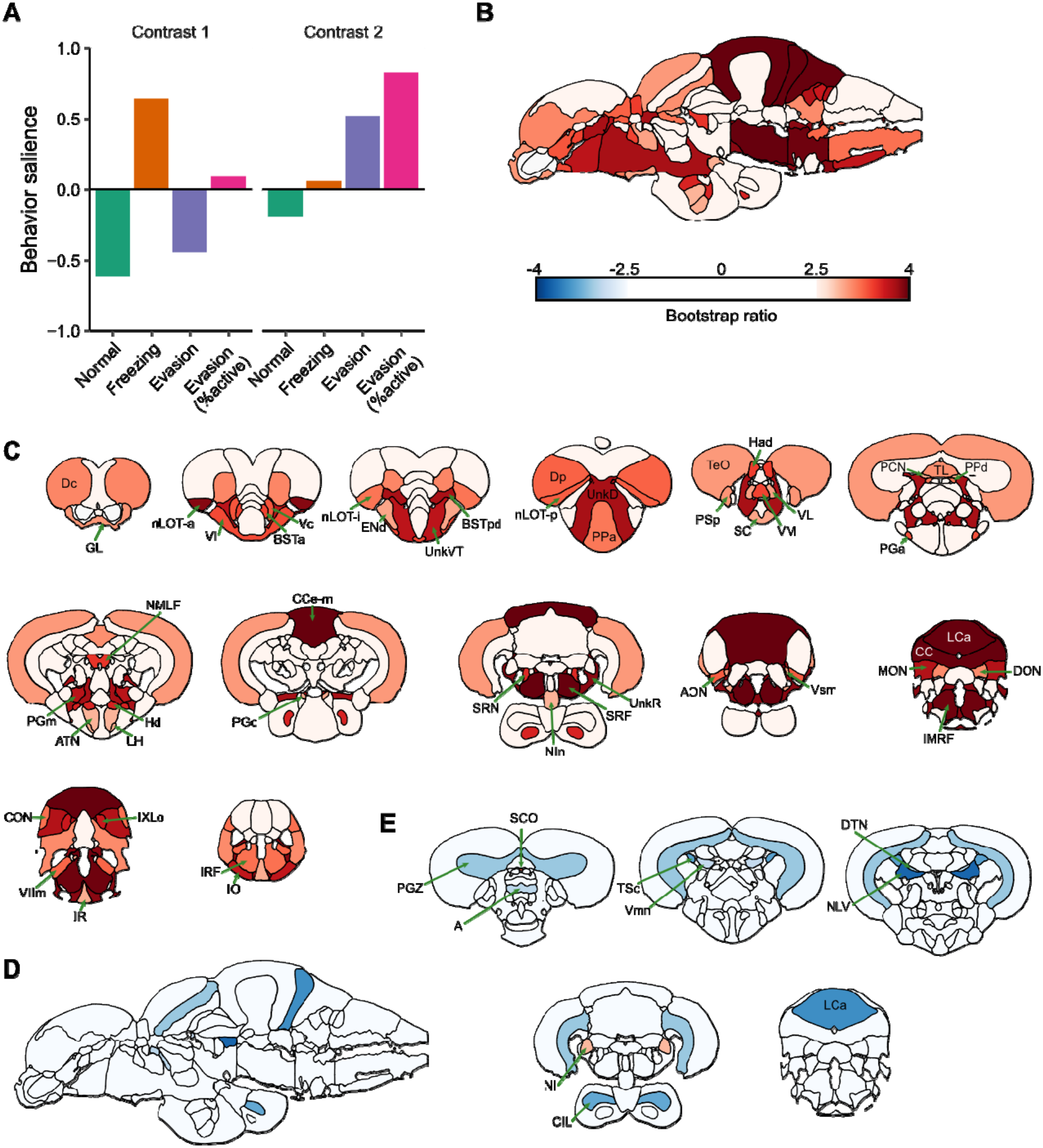
Partial least squares (PLS) analysis to identify brain regions that covary with behavior. A) Top two contrasts and related behavioral saliences identified from PLS analysis. B) Bootstrap ratios of brain saliences for the first contrast on a sagittal image of the adult zebrafish brain. Only bootstrap ratios beyond ± 2.5 are depicted. C) Coronal slices with same coloration as part B. D) Bootstrap ratios of brain saliences for the second contrast on a sagittal image. E) Coronal slices of bootstrap ratios correspond to the second contrast. Brain region abbreviations can be found in Table S6-1. Grayed out areas represent white-matter tracts; N = 87.

To determine which brain regions most consistently contributed to the behavioral contrasts, we calculated bootstrap ratios (BSRs, i.e., salience scores). BSRs capture a combination of the magnitude and stability of regional contributions to a contrast (McIntosh and Lobaugh, 2004). For the first contrast (**Figures 6B** and **C****, S6-1A**), we found numerous positive contributions [BSRs > 2.5; i.e., neural activity covaried with freezing]. Although PLS is multivariate, we have provided example univariate relationships between *cfos* activity and behavioral contrasts to give an intuition as to what relationships are being captured (**Figure S6-2**). This widespread positive activity pattern suggests that freezing behavior, despite the lack of movement, is not passive at the neural level but instead involves coordinated brain-wide activity. The cerebellum in particular stands out with several regions having very high BSRs: the CC (cerebellar crest; BSR = 4.7), CCe-m (molecular layer of the corpus cerebelli; BSR = 4.0), MON (medial octavolateralis nucleus; BSR = 3.6), and LCa (caudal lobe of the cerebellum; BSR = 6.5). The MON is one of the primary sites of lateral line input to the brain (Valera et al., 2021; Wullimann and Grothe, 2013) and interacts with the rest of the cerebellum (Dohaku et al., 2019; Pose-Méndez et al., 2023). Thus, this data suggests that freezing involves constant monitoring for movement in the water, possibly to maintain posture and minimize drifting. Maintenance of freezing is likely driven by regions of the reticular formation, which projects to the spinal musculature (Chong and Drapeau, 2007; Huang et al., 2013), where we also see strong associations with contrast 1: the SRF (superior reticular formation; BSR = 5.0), IMRF (intermediate reticular formation; BSR = 4.0), and SRN (superior reticular nucleus; BSR = 3.4). Finally, several regions of the telencephalon positively covaried with contrast 1, such as the nuclei of the lateral olfactory tract (nLOTa: BSR = 3.8, nLOTp: BSR = 3.0), parts of the bed nucleus of the stria terminalis (BSTpd: BSR = 3.8, BSTa: BSR = 3.22), the central and lateral portions of the ventral telencephalon (Vc: BSR = 3.1; Vl: BSR = 3.2), and the posterior and central portions of the dorsal telencephalon (Dp: BSR = 3.0; Dc: BSR = 2.8). These regions are likely working to scan and evaluate external chemosensory threats where the nLOTs and Dp receive input from the olfactory bulbs (von Bartheld et al., 1984; Yáñez et al., 2022). These regions may work in concert with the BSTs, which are a part of the extended amygdala, to assign valence to chemosensory stimuli (Porter and Mueller, 2020). The BSTs likely interact with the Vc, which is proposed to be a part of the central amygdala (Porter and Mueller, 2020), a region important for the expression of emotional learning like fear conditioning (Ciocchi et al., 2010; Wilensky et al., 2006). Finally, the Dc is thought to be homologous to mammalian iscortex (Mueller, 2022; Mueller et al., 2011), integrating input from parts of the brain and projecting to midbrain regions to organize brain-wide activity (Yáñez et al., 2022).

For the second contrast (**Figures 6D**, **6E** and **S6-1B**), there were only two regions with positive BSRs, covarying with evasive behavior: the nucleus isthmi (NI: BSR = 2.6) and the subcomissural organ (SC: BSR = 3.1). The NI is important for visual attention and modulating retinotectal input, particularly in response to looming stimuli that signal danger (Fernandes et al., 2021; Gruberg et al., 2006; Henriques et al., 2019). Thus, fish with more evasive behavior may be driven by greater attentiveness to potential visual threats. The SCO has recently been found to secrete glycoproteins that form Reissner fiber’s in the spinal cord that help detect spinal curvature (Cantaut-Belarif et al., 2018; Orts-Del’Immagine et al., 2020), and so this region may be more active in order to repair Reissner fibers that come under stress due to the rapid changes in spinal curvature that occur during evasive movements. Regions related to foraging behavior and mechanosensory feedback negatively covaried with contrast 2 and thus are shut down during evasion and more engaged during normal swimming. For example, we have negative BSRs for regions important for spatial navigation like the periventricular gray zone of the optic tectum (PGZ: BSR = −2.8) and the anterior thalamic nucleus (A: BSR = −2.5), which are involved in processing a wide range of visual information (Baier and Scott, 2024; Baier and Wullimann, 2021; Cheng et al., 2017) and the dorsal tegmental area (DTN: BSR = −2.7), which is critical for monitoring head direction (Petrucco et al., 2023); all of these would be important for foraging behavior. The central nucleus of the inferior lobe (CIL: BSR = −3.0) is part of a region implicated in feeding behavior (Muto et al., 2017; Wee et al., 2019), but may also play a role in integrating sensory input (Bloch et al., 2019). Finally, several regions involved in sensory feedback are shut down during evasive behavior, likely in anticipation of the large inputs that would occur during rapid movements. These include regions important for auditory processing like the nucleus of the lateral lemniscus (NLL: BSR = −3.1) and central nucleus of the semicircular torus (TSc: BSR = −3.0) (Yáñez et al., 2024), and vestibular information like the nucleus lateralis valvulae (NLV: BSR = −3.6), and the caudal lobe of the cerebellum (LCa: BSR = −2.9) (Pose-Méndez et al., 2023).

### Brain networks underlying individual differences in behavior

The brain is best thought of as a network of interacting regions whose complex interactions give rise to behavior. To determine the patterns of activity that are present during different types of fear memory expression, we generated networks from correlated *cfos* activity between brain regions for each experimental group (**Figure 7**). The overall pattern of correlations (**Figures 7A** & **B**) is similar in the groups exposed to the behavioral tank (i.e., non-reactive, evading freezer, and freezer groups; similarity of matrices: r’s=0.41-0.43) while clearly differing from the habituation group that was not exposed to the behavioral tank (r’s=0.18-0.24). To identify which regional interconnections were significant for network generation, we applied a threshold (FDR ≤ 0.001; **Figure 7C**). This identified fewer significant correlations in the non-reactive group (306) than the evading freezer (588) and freezer groups (655), suggesting the freezer groups form a more coherent brain network.

**Figure 7.**
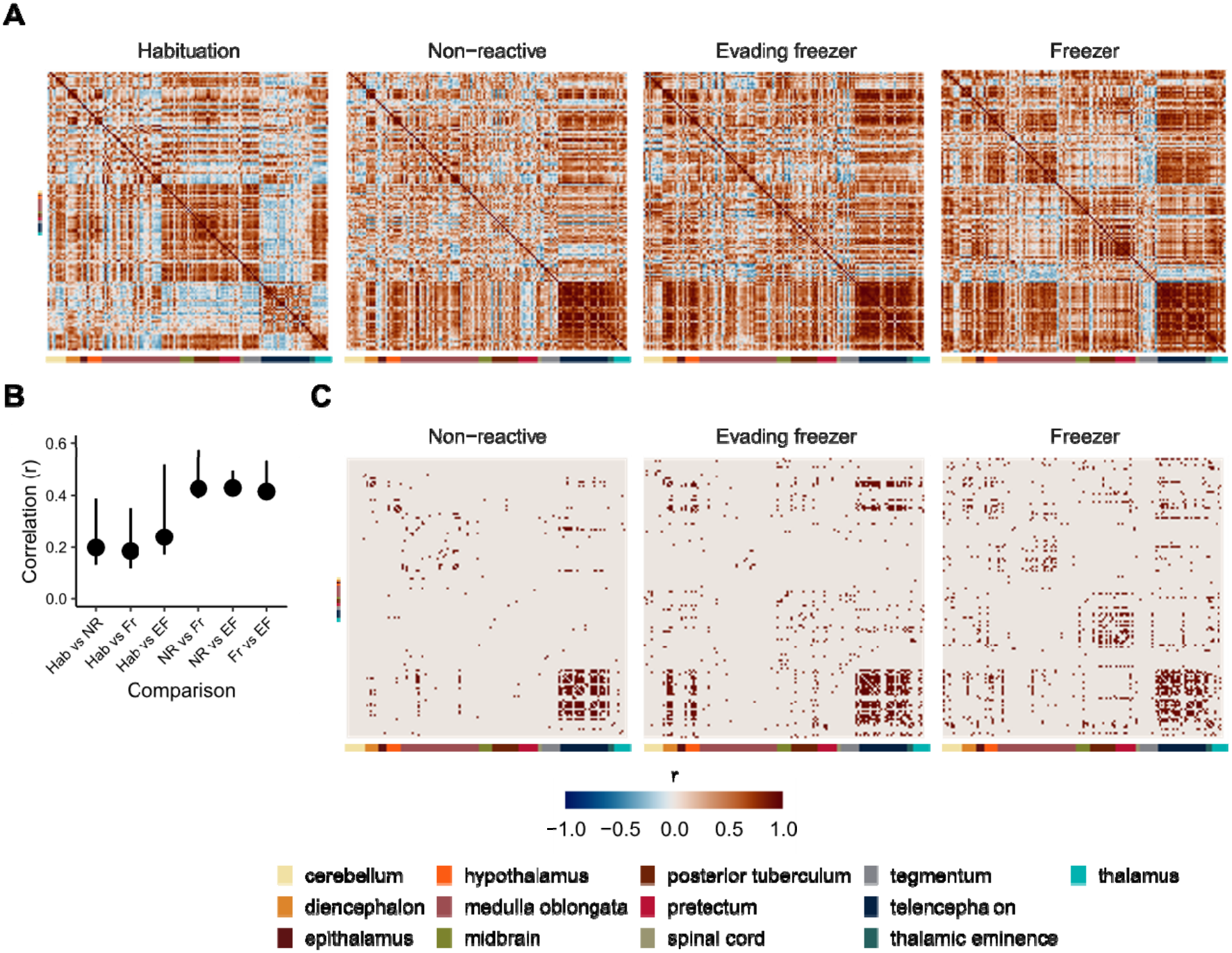
Covariation between brain regions of different behavioral groups. A) Correlation matrices of cfos activity between brain regions. Each row/column is an individual brain region, and each entry is the Pearson correlation of cfos activity across animals in a group. Colored bars represent the ontological level of each brain region. B) Pearson’s correlations of matrices from each group with one another. Error bars are bootstrapped 95% confidence intervals. C) Correlation matrices with only suprathreshold correlations (FDR ≤ 0.001). Habituation: n = 19; non-reactive, n = 17; evading freezer: n = 16; freezer: n = 21.

To generate insight into how the patterns of brain activity in the evading freezer and freezer groups differed, we generated functional networks (**Figure 8**). In these networks, the nodes represent brain regions and edges represent the presence of a suprathreshold correlation (**Figure 7C**). To better visualize how the networks in these two behavioral groups differed, we identified edges whose confidence intervals did not overlap with those of the other behavioral network or the non-reactive network (**Figure 8A**, solid lines). Significant edges/correlations that had overlapping confidence intervals are depicted as dashed lines.

**Figure 8.**
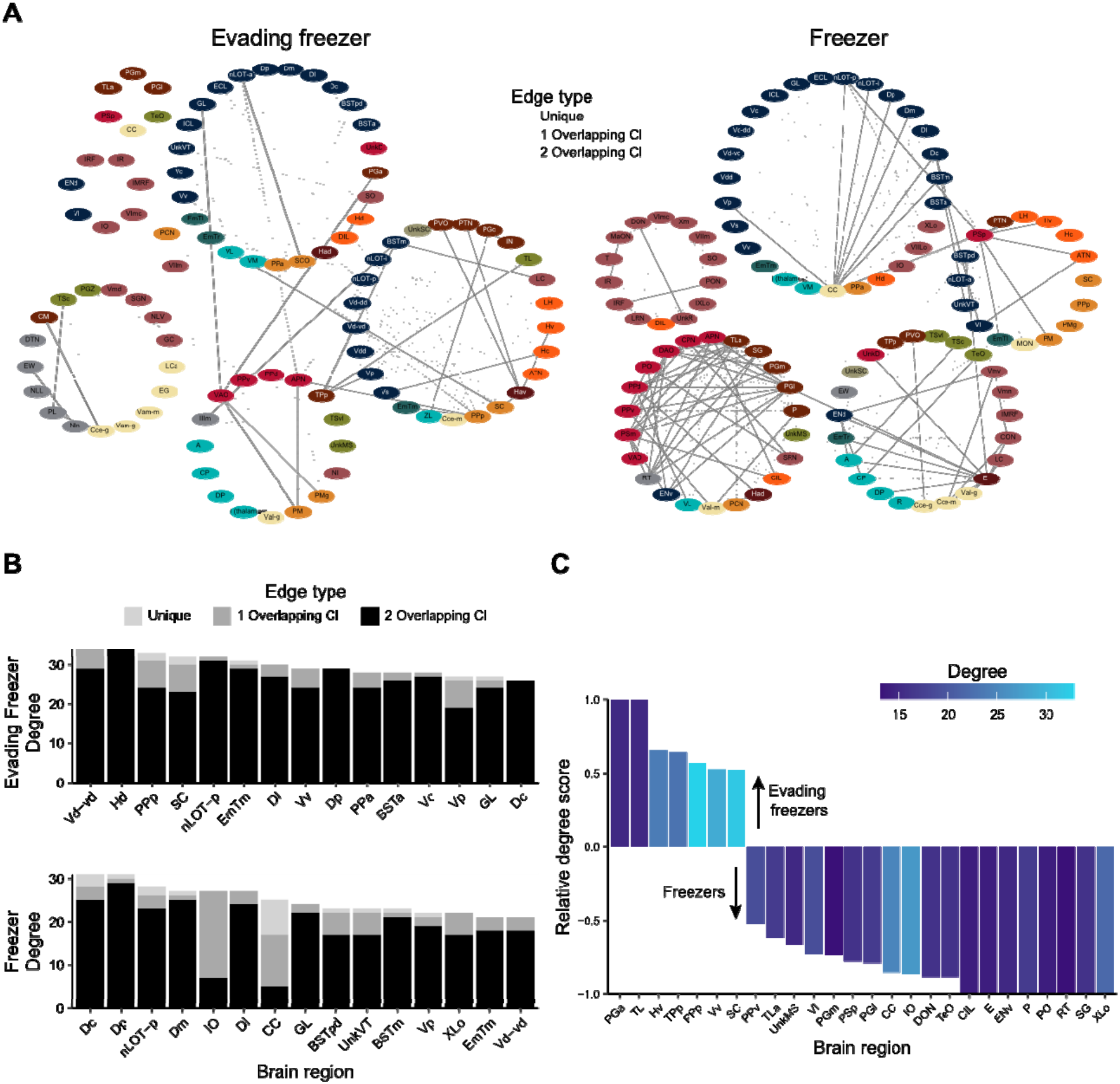
Functional brain networks underlying fear memory expression. A) Functional networks where nodes represent brain regions and edges the presence of suprathreshold correlations (FDR ≤ 0.001). Node colors correspond to ontological levels from figure 7. Solid lines are edges whose 95% confidence intervals do not overlap with either of the other two networks (evading freezer, freezer, or non-reactive). Dashed lines represent edges that have 95% confidence intervals that overlap with one (dark gray) or both (light gray) of the other networks. B) Degree centrality of the top 15 nodes. Bars are colored according to the number of edges that arise from unique (light gray) or overlapping confidence interval (dark gray = 1 overlap or black = 2 overlaps). Overlaps defined the same as in part A. C) The relative degree score for nodes enriched in the evading freezer (positive) or freezer (negative) functional networks. Only those nodes with a relative degree score of more than ± 0.5 and degree greater than 12 are shown. Brain region abbreviations can be found in Table S6-1.

All networks (evading freezer, freezer, non-reactive, and vehicle treated fish; **Figures 8A and S8-1, S8-2**) have strong interconnectivity in the telencephalon (blue), including high degree centrality in both the lateral zone of the dorsal telencephalon [Dl (degree: evading freezers = 30, freezers = 27, non-reactive = 25, and vehicle = 23)] and the medial zone [Dm (degree: evading freezers = 23, freezers = 27, non-reactive = 20, and vehicle = 27)]. These regions are notable for being the putative homologs to the mammalian hippocampus (Dl) and basolateral amygdala (Dm) (Anneser et al., 2024; Ganz et al., 2012; Mueller et al., 2011; Porter and Mueller, 2020), both of which are well known to be critical for contextual fear conditioning in mammals (Tovote et al., 2015). This finding is particularly interesting because it suggests that these regions are activated non-specifically simply by being placed back into the conditioning tanks. High inter-connectivity of these regions does not occur following an initial exposure to a novel tank (Rajput et al., 2025), and thus this pattern of activity may represent a general response to being placed back into a previously experienced context.

The evading freezer network is notable for its numerous interactions between the subpallium (ventral telencephalon) and hypothalamus (orange). In particular, there is high degree centrality in the mammalian homologs of the central and medial amygdala (Vd-vd, Vc, and Vp), striatum (Vd), and septum (Vv) (Porter and Mueller, 2020; Rink and Wullimann, 2004). Thus, in evading freezers it appears that the Vd-vd, Vc, and Vp may help evaluate the threat while vacillation between active and passive responses is likely regulated by the septal homolog (Vv; Besnard et al., 2019; Rizzi-Wise and Wang, 2021) with the evasive behavior program likely initiated by the striatal homolog (Vd). This interpretation is supported by the fact that the Vv is less engaged in the freezer network (degree = 9) where there is less switching between freezing and evasive behaviors (**Figures 4D & F**). The Vv also has extensive projections to the preoptic area and hypothalamus (Rink and Wullimann, 2004; Rizzi-Wise and Wang, 2021), which contains several regions overrepresented in the evading freezer group [i.e., the dorsal and ventral hypothalamus (Hd and Hv) and the anterior and posterior preoptic areas (PPp and PPa)]. These preoptic and hypothalamic regions are key regulators of autonomic functions, homologous to their mammalian counterparts (Biran et al., 2015). This suggests that the evading freezer behavioral type may be driven by an elevated stress response relative to freezers.

In contrast, the freezer functional network is notable for interactions between the telencephalon and the cerebellum, particularly the CC which has several unique connections with the dorsal telencephalon and has much higher degree than in the other two networks (CC degree: freezers = 25, evading freezers = 2, non-reactive = 0, vehicle = 1). The other notable aspect of the freezer network is the cluster that contains several unique connections between the pretectum and posterior tuberculum. Several parts of the posterior tuberculum stand out as being highly interconnected in freezers, like the medial and lateral preglomerular nuclei (PGm and PGl, respectively), and the posterior thalamic nucleus (P). These nuclei are thought to serve as key sensory processing regions for vision, gustation, and olfaction that then project to the pallium, not dissimilar in function to the mammalian thalamus (Bloch et al., 2020; Mueller, 2012; Northcutt, 2006). Thus, in freezers, enhanced sensitivity to sensory input may be modulating the pallium to signal to the cerebellum to control and maintain the freezing posture and suppress evasive behavior.

## Discussion

The present study is the first to identify the presence of individual differences in fear memory expression in zebrafish. We built on prior work that found fish exhibit a mix of active and passive responses to CAS (Hall and Suboski, 1995; Maximino et al., 2018; Speedie and Gerlai, 2008; von Frisch, 1938). We can now add that individuals express fear memories in different ways: some individuals have high levels of freezing interspersed with normal swimming (i.e., freezers) whereas others intersperse freezing with evasive behavior (i.e., evading freezers). Finally, we uncovered patterns of brain activity that underlie these behavioral types, finding that engagement of the telencephalon tends to be common across behavioral types, whereas differences arise in connections to the cerebellum, preglomerular nuclei and pretectum in freezers, and preoptic area and hypothalamus in evading freezers.

We found that fish respond to CAS with an increase in both freezing and evasive behaviors, with freezing being the more robust and persistent response. These responses to CAS are consistent with previous reports of active and passive responses in fish (Hall and Suboski, 1995; Maximino et al., 2018; Speedie and Gerlai, 2008; von Frisch, 1938). We also found that strain had a modest effect on behavior, with the most prominent finding being that TU fish were overrepresented in the non-reactive cluster during fear memory expression. This echoes prior work where the TU strain was found to have a faster rate of contextual fear memory extinction than the AB and TL strains (Kenney et al., 2017). This could be due to a reduction in learning, as TU fish did not increase their freezing response to CAS as much as the other strains (**Figure 2A**). We also saw a sex difference with respect to evasive behaviors, where males are more likely to increase their evasive behaviors than females when CAS is present. Freezing was largely similar across sexes. This is consistent with a recent study that found both male and female fish increase freezing in response to CAS (Akinrinade et al., 2023). However, the prior study found no change in evasive behaviors CAS and no sex differences. We suspect this negative finding may be because the prior work examined overall evasive behaviors, which tend to be overwhelmed by high levels of freezing. We handled this issue by assessing evasive behavior as a percentage of time not freezing, giving us greater sensitivity to uncover subtle behavioral differences.

A key finding from the present study was the identification of individual differences in how fish behave during fear memory recall. We found that fish behavior falls into one of four groups: non-reactive, evaders, evading freezers, and freezers. Groups high in freezing (evading freezers and freezers) were the most clearly associated with responses to CAS and fear memory. Differentiating fish with high levels of freezing into evading freezers from freezers, instead of collapsing them into a single group, is supported by several lines of evidence: (i) Division into evading freezers and freezers provides the most optimal clustering that is robust to parameter changes while maximizing internal clustering metrics (**Figure 4 and Figure S4-1**). (ii) Transitions between behavioral states in freezers and evading freezers are distinct (**Figure 4D-F**). This finding is important because behavioral transitions are determined from second-to-second data that were not used for clustering. These data suggest that, despite relatively small differences in overall levels of evasion in evading freezers and freezers, they organize their behavior differently (**Figure 4D-F; Figure S4-3**). In particular, compared to freezers, evading freezers have an increase in evasive bout length (**Figure 4E**) and drastic increase in the likelihood of transitioning from freezing to evasion (**Figure 4F**). (iii) The differences in the patterns of brain activity in evading freezers and freezers (**Figure 8**) is consistent with them being distinct behavioral types (discussed in more detail below). We do not consider evaders a fear-behavior phenotype because a substantial number of animals fell into the evader category in the vehicle treated fish and the CAS treated fish during pre-exposure (**Figure 5A**). This suggests that an increase in evasion in the absence of freezing may be a behavioral response to netting and placement in the tank rather than to the alarm substance (Shishis et al., 2023). We suspect this elevated evasive behavior may also be due to the relatively small size of the tank used in this study, which we chose to limit the amount of CAS needed to elicit a fear response.

Our findings on individual differences in fear memory expression are distinct from prior work examining such differences in rodents and fish that only use freezing levels to classify animals as either high or low in fear expression. For example, using an active avoidance paradigm with CAS, Maximino and colleagues (2018) classified zebrafish as high or low in avoidance during memory expression, finding that high avoiders had higher levels of freezing and lower levels of erratic movements during CAS exposure. A similar approach has been taken in rats, where individual differences have been identified by differentiating animals based on their freezing responses to conditioned stimuli (Graham and Richardson, 2016; Walters et al., 2016). Interestingly, rats have been found to exhibit a mix of active (darting) and passive (freezing) fear behaviors during fear conditioning to an electric shock where females exhibit more darting than males (Gruene et al., 2015; Mitchell et al., 2022). Elevated darting was also associated with less freezing in rats. These findings mirror some of the findings in the present study, such as the presence of both active (evasive) and passive (freezing) fear behaviors. However, we do not find as clear of a sex difference in active and passive behaviors during fear memory expression, and we find that evasive behaviors can coexist with high levels of freezing. These differences may be due to the use of a different unconditioned stimulus (CAS versus electric shock) and/or species.

We find that the cerebellum stands out as being strongly associated with freezing behavior. This conclusion is supported by both the PLS and network analysis. PLS analysis found that neural activity of several cerebellar regions (CC, CCe-m, LCa, and MON) strongly contributed to the covariation of brain activity with freezing behavior. The network analysis also identified the CC as a hub with many unique interactions with the pallium and subpallium. Relatedly, the inferior olive (IO) was also identified as important by the PLS and network analysis. The IO is a key source of instructive input to the Purkinje cells of the cerebellum that process information and serve as its primary output (De Zeeuw et al., 1998; Dohaku et al., 2019; Pose-Méndez et al., 2023). A role for the cerebellum in fear-induced freezing is consistent with prior work in goldfish (Yoshida et al., 2004), and recent views of the cerebellum playing an important role in socio-emotional behavior alongside its more well established involvement in motor coordination (Strata, 2015). In looking closer at the functional connectivity of the cerebellum in freezers, we find connections between the CC, IO, and Dm where the Dm is thought to be the teleostan homolog of the basolateral amygdala (Mueller, 2022; Porter and Mueller, 2020) which has been implicated in fear memory learning in both zebrafish and rodents (Fanselow and LeDoux, 1999; Lal et al., 2018; Sun et al., 2020-1-28). Thus, freezing here may be driven by an amygdala-cerebellar circuit (Rudolph et al., 2023).

Another collection of brain regions that stands out as key to freezing behavior are the posterior tuberculum and pretectum. Within the posterior tuberculum, the preglomerular nuclei are amongst the top contributors to freezing in the PLS and network analysis (particularly the PGm which is supported by both sets of analyses). Preglomerular nuclei receive a variety of sensory inputs and project to the pallium, making it functionally analogous to the mammalian dorsal thalamus (Mueller, 2012; Trinh et al., 2026; Wullimann, 2020). In particular, the PGm in goldfish, a related teleost species, receives chemosensory input (Northcutt, 2006). Thus, the PGm may work together with other chemosensory regions to recall the CAS memory, like the nLOTa, nLOTp, and Dp that positively co-vary with freezing behavior. Pretectal nuclei, like the PSp and PPv, were also identified as important by both the PLS and network analysis. The pretectum is key to visual processing as these regions receive retinal input and have reciprocal interactions with the optic tectum (Yáñez et al., 2018). Interestingly, several pretectal nuclei, including the PPv, receive or send projections to the CC of the cerebellum (Yáñez et al., 2018). Taken together, these data suggest that the freezing behavior in the absence of elevated evasive behavior, may be driven by a bias towards generating memories from combined visual and chemosensory inputs. These sensory memories then engage a cerebellar-telencephalic circuit to initiate the coordination of freezing behavior. However, further anatomical and functional manipulations would be required to more directly draw these conclusions.

In the functional brain network of evading freezers, we see greater engagement of the hypothalamus and preoptic areas than in freezers. For example, the network analysis identified the SC, PPp, and PPa, parts of the preoptic area, as high in degree and for having a higher proportion of unique edges than other brain regions. These regions formed a cluster that has many edges to hypothalamic and subpallial regions (**Figure 8A**). The preoptic area of zebrafish has been found to be a neurosecretory area, roughly homologous to the paraventricular nucleus in mammals (Herget et al., 2014). This region, along with other hypothalamic nuclei, modulate a broad spectrum of behaviors and physiological processes, including the hypothalamic-pituitary-interrenal (HPI) axis that regulates the release of cortisol, a central stress hormone (Biran et al., 2015; Wendelaar Bonga, 1997). Relatedly, CAS has been found to increase cortisol in zebrafish (Abreu et al., 2017); and in the frillfin goby (*Bathygobius soporator*) cortisol modulates the behavioral effect of CAS (Barreto et al., 2014). Thus, it may be the case that individual variation in the status of the HPI axis in zebrafish may determine if zebrafish exhibit a combination of evasive and freezing behaviors (i.e., evading freezers) or freezing interspersed with normal swimming (i.e., freezers).

In conclusion, we have identified the presence of individual behavioral variation in how zebrafish express fear memories. We find that the strain and sex of fish affect fear responses and that the behavioral response during learning is predictive of memory expression. Different types of fear memory expression are associated with differences in the activity of brain regions like the cerebellum, preglomerular nuclei, preoptic area and hypothalamus. Future work will be needed to probe the function of these circuits and determine how they may be differentially wired to give rise to variation in behavior.

## Materials and Methods

### Subjects

Animals were female and male zebrafish from the AB, TU, TL and WIK strains 6 to 9 months of age. Fish were bred in-house and raised at Wayne State University and were within two generations of fish obtained from the Zebrafish International Resource Center (ZIRC - University of Oregon). Fish were housed in a re-circulating system (Aquarius Fish System, Aquatic Enterprises) with a 14:10-hour light-dark cycle (lights on at 8:00 a.m.; pH 7.5 ± 0.2 °C; water conductivity 500 ± 10 μS; temperature 27.5 ± 0.5 C). Fish were fed two times a day with dry feed in the morning (Gemma 300, Skretting) and brine shrimp in the afternoon (Artemia salina; Brine Shrimp Direct). All the procedures were approved by the Wayne State University Institutional Animal Care and Use Committee (Protocol ID: 21-02-3238).

For the behavioral experiments, animals were housed in male/female pairs (2 female/male pairs per tank separated by a transparent divider) in a 2L tank for one week before behavioral manipulations. The pair-housing system minimizes potential stress induced by social isolation (Parker et al., 2012), and maintains a consistent environment while allowing for the identification of individual fish across days. Behavioral experiments were performed between 11:00 and 15:00 h. Fish were moved to the behavioral room 1 hour before experiments to allow animals to adapt to the new environment. The fear conditioning task consisted of 3 days: day 1: habituation; day 2: conditioning; day 3: memory recall (**Figure 1A**). After behavioral recording, animals remained in the room for 1 hour before being moved back to the re-circulating system. Sex was determined using three secondary sex characteristics: body shape, color, and presence of tubercles on the pectoral fins (McMillan et al., 2015). At the end of the experiment, we confirmed sex after euthanization by dissection to identify the presence of eggs. To ensure data reliability, eight independent batches of offspring for each strain were used for behavioral testing (n = 12 – 16 per strain in each batch). All behavioral testing was carried out using a counterbalanced design across sex and placement in tank (front/back).

### Fear conditioning

The fear conditioning protocol was adapted from prior work (Kenney et al., 2017) to use the Zantiks automated system (Zantiks Ltd) and conspecific alarm substance (CAS) as the unconditioned stimulus (Maximino et al., 2018). Two tanks (20 x 7 x 9 cm) were recorded simultaneously during the experiments at 60 fps. A pilot study was performed to determine the best CAS concentration starting with previous work in which a concentration of 3.5mL/L elicited fear-related responses up to 7 days after exposure (Maximino et al., 2018). We found a 1:10 dilution of this dosage (0.35 mL/L) elicited a robust freezing response with some variation on memory day (**Figure S2-1**). Fish were tracked using DeepLabCut (Mathis et al., 2018) by labelling three points (head, trunk and tail). We used ResNet101 network and trained a deep neural network on 200 manually labelled frames (divided across the four strains and both sexes), the network was then refined and improved by correcting outliers to include 578 additional frames.

### CAS extraction

CAS is a substance released by fish when their epithelial cells are damaged, typically in response to a strike from a predator. We extracted CAS as previously described (Egan et al., 2009; Quadros et al., 2016). Briefly, CAS was obtained from donor fish that were first euthanized via rapid cooling in 4 °C water. Fish were placed in a 10 cm petri dish and epidermal cells were damaged via 10 shallow incisions on both sides of the body using a surgical blade on ice without drawing blood. 10 mL of distilled water was then added to the dish and it was gently agitated to ensure complete coverage of the lacerated areas on the fish body. Because 4 different strains were used for the behavioral experiments, we prepared stock solutions of alarm substance by mixing extract from 1 fish per sex/strain and then aliquoting before storing at −20 °C for no longer than 4 weeks.

### Behavioral analysis

To automate capture of behavior we developed a random forest machine learning model. After fish were tracked using DeepLabCut, we calculated postural information for each individual, extracting 34 parameters (**Table S1-1**) over a sliding window of 750 ms (45 frames). We manually labelled 14,199 frames per behavior: (1) freezing, the absence of movement with fast opercular movements, (2) erratic movements, zig-zagging movement of at least 3 quick C-bends, (3) burst swimming, a rapid increase of speed in one direction, 4) straight swimming, normal speed swimming in one direction and, (5) normal turns, (**Figure 1B**). We used an 80/20 (train/test) split for training and testing the model. Optimization of the model was done by testing a range of variables per level (1-15) and trees. We found that 10 variables and 2,000 trees yielded the lowest out of bag error rate (2.88%). Results of testing on 20% of unseen data also yielded excellent results with errors of no more than 6% and typically much lower (**Figure 1C**).

To analyze the temporal organization of zebrafish behavior, we modeled behavioral state transitions using discrete-time Markov chains. Behavioral sequences were determined by calculating the modal behavior in 750 ms time windows. This time window matched the length of the sliding window used in the random forest classifier. Transition matrices were calculated using maximum likelihood estimation using the markovchain package (version 0.9.5) in R (Spedicato, 2017). Matrices were fit using data pooled across all animals within each experimental group and day. Ninety-five percent confidence intervals were computed and used for determining significance.

### Identification of behavioral clusters

Clusters were identified by creating a k-nearest neighbor network and then applying the Louvain community detection algorithm (Blondel et al., 2008). We only used behavior on memory day from fish exposed to CAS. To calculate distances for the k-nearest neighbor network, we first standardized the behavioral data (percent freezing, percent evasion, and percent evasion as a proportion of active time). We then calculated similarity scores between individual fish:

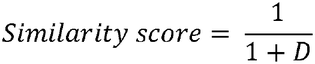

where D is the Euclidean distance between each fish in the three-dimensional behavioral space. We selected the optimal k value by calculating internal clustering metrics using the clusterCrit package (version 1.3.0) at each value of k: Calinski–Harabsz index (Caliński and Harabasz, 1974), silhouette index (Rousseeuw, 1987), and Davies–Bouldin index (Davies and Bouldin, 1979). Based on these data, we chose k = 74, which fell within a range that optimized internal clustering metrics and showed robustness to small changes in k (**Figure S4-1**). Clustered data were visualized using a uniform manifold approximation (UMAP) (McInnes et al., 2020) as implemented in the UMAP R package (version 0.2.10).

To identify clusters in new data, such as during pre-exposure and exposure or in fish used for brain imaging analysis, we first standardized the new data using the parameters from the initial clustering described above. We then assigned clusters to each fish based on the number of connections to its 33 nearest neighbors in the initial network. Thirty-three was chosen because it is half the size of the smallest cluster identified.

We also clustered the data using two other approaches: using only two variables (percent freezing and percent evasion) and by generating isometric log ratios. The latter approach was motived by the fact that the data is compositional (i.e., percent freezing + percent evasion + percent normal swimming = 100%) (Aitchison, 1982; Greenacre, 2018). We generated an orthonormal basis using an isometric log ratio transformation (Egozcue et al., 2003):

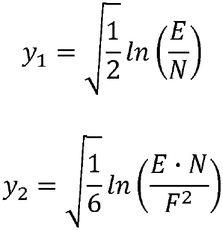

Where E = percent evasion, N = percent normal swimming, and F = percent freezing. Prior to the transformation, zeros in the data were replaced with 0.01% (about half the minimum non-zero behavioral percentage measured). These variables were then used for clustering as described above.

### Tissue clearing and staining

Brains were collected 15 min after the end of fear memory expression. Fish were rapidly euthanized via submersion in ice cold water and heads were removed and fixed in 4% paraformaldehyde overnight. Animals that were assigned the wrong sex (determined post-mortem as described above) were removed from data analysis, as well as its paired fish (< 2%). After overnight fixation, brains were carefully dissected in cold PBS. Brains damaged during the dissections were discarded. Tissue staining and clearing was performed using iDISCO+ (Renier et al., 2016) as previously described (Rajput et al., 2025). After dissections, samples were washed three times with PBS (30 min washes) at room temperature followed by dehydration using methanol/water (20, 40, 60, 80 and 100% methanol for 30 min each). Samples were placed in 100% methanol for one hour and chilled in the fridge (4° C). They were then incubated at 4° C overnight in 5% hydrogen peroxide in methanol. The next day, samples were rehydrated in methanol/water (80, 60, 40 and 20%, 30 min each) followed by two PBS washes, one PTx.2 (PBS with 0.2% TritonX-100) of one hour each and an overnight incubation in 5xSSCT buffer (sodium chloride sodium citrate and 0.2% TritonX-100). The following day, an acetylation step was performed with 0.25% v/v acetic anhydride for 30 min followed by three washes with ultrapure water for 5 min each. Samples were then incubated in hybridization buffer (Molecular Instruments) for 15 min at room temperature followed by one hour incubation at 37° C with the same buffer. A solution of hybridization buffer and *cfos* probes (2 pmol per sample) was then prepared and samples were incubated with the probes for 3 days. Samples were then washed three times using probe wash buffer (Molecular Instruments) followed by two 5xSSCT washes of one hour each. Finally, samples were incubated for 1 h with amplification buffer (5×SSC, 0.1% Tween 20, 10% dextran sulfate) and then incubated in the dark for 3 days in 125 µL of amplification buffer and 2.5 µL of each B1 hairpin (7.5 pmol). After amplification, samples were kept in the dark and washed five times in 5xSSCT buffer and left overnight. Clearing then began with dehydration in methanol/water mixtures (20, 40, 60, 80, 100, and 100%) for one hour each, followed by three hours incubation in dichloromethane/methanol (66%/33%) solution. Samples were then incubated two times for 15 min with dichloromethane. Lastly, the solution from the tubes was removed and dibenzyl ether was added to the samples in which they remained until imaging.

### Whole-brain imaging

Cleared samples were imaged via light-sheet microscopy on an UltraMicroscope II (Miltenyi Biotec). Samples were immersed in dibenzyl ether and affixed to a pedestal using a refractive index matched ultraviolet cured resin (NOA 61, Norland Products). Images were acquired at 5× magnification. Illumination of samples was from the right laser using dynamic horizontal focusing and a step size of 4 µm. Autofluorescence images were acquired using a 480 nm laser and used for registration. C*fos* images were acquired using an illumination laser of 641 nm. The exposure time was 200 ms, sheet width of 50%, and a sheet numerical aperture of 0.156. Images were acquired as 1 × 2 tiles and stitched using TeraStitcher (Bria and Iannello, 2012).

### Automated cell detection

Detection of *cfos* positive cells was done using CellFinder, version 0.4.20 (Tyson et al., 2021) as previously described (Rajput et al., 2025). Briefly, cell candidate detection was first performed and those cell candidates that were closer than 9 µm to one another were removed. We then applied the CellFinder artificial neural net supervised machine learning algorithm to differentiate cells from non-cells. To train the CellFinder network, we labelled 25,460 cells and non-cells and trained the network to an accuracy of 97.7%. For one brain, cell counting was unsuccessful in the telencephalon and so these regions were excluded from the analysis.

### Registration to AZBA

Images were registered to the adult zebrafish brain atlas (AZBA; Kenney et al., 2021) using advanced normalization tools (version 2.4.4; Avants et al., 2011) to automatically parcellate brains into individual brain regions as previously described (Rajput et al., 2025). We initially generated a template from autofluorescence images that was a combination of a previously generated template (Rajput et al., 2025) and 10 brains from the present study. The autofluorescence image from AZBA was then registered to the template and the transformation from each individual brain to the template was used to bring the AZBA autofluorescence image into the space of individual brains. We had poor registration of the olfactory bulbs in three brains and so these regions were excluded from the analysis for these samples.

### Partial least squares (PLS) analysis

We implemented PLS analysis (Krishnan et al., 2011) in R to identify covariation between behavior and regional brain activity. Because of batch-to-batch variation in the brightness of *cfos* images, cell counts were normalized based on the median overall *cfos* count for each cohort. This normalized data was then used for PLS (n = 87). Only fish exposed to CAS during training day were included in the analysis. Using mean-centered behavioral PLS correlation, we applied a singular value decomposition to a matrix of correlations between normalized *cfos* counts and four behaviors: percent normal swimming, percent freezing, percent evasive swimming, and percent evasive swimming while active (i.e., not freezing). Because the behavioral data is measured as percentages, some of which were skewed, we used an arcsine square root transformation prior to the PLS analysis:

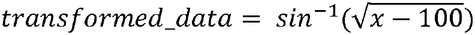

To determine which latent variables to consider, we applied permutation tests (resampled 10,000 times without replacement) and used this to calculate p-values associated with each singular value. To determine the reliability of brain region contributions to a specific latent variable, we used bootstrapping (resampled 10,000 times with replacement). Bootstrap ratios were calculated by dividing the salience associated with each brain region by its standard error calculated from bootstrapping. Bootstrap ratios are similar to z-scores and thus we consider a threshold of ± 2.5 as significant, which is roughly equivalent to a two-tailed p-value of 0.01.

### Brain network analysis

Brain networks were derived from Pearson correlation matrices capturing covariation between *cfos* counts per brain region across animals in each experimental group. Matrices were thresholded using a false discovery rate (FDR) ≤ 0.001 (Benjamini and Hochberg, 1995). For calculating the initial p-values from Pearson’s correlations, we used the lowest number of observations (n = 16) from the groups used for making networks. We made this conservative correction because the p-value is highly sensitive to the number of observations, which would cause a large skew in the number of significant edges identified for each group. To capture edges that were unique to each network, we identified edges whose 95% confidence intervals did not overlap between networks. Only those edges that were significant (i.e., FDR ≤ 0.001) and did not have overlapping confidence intervals (CI) were labelled as ‘unique’. Those with only one overlap were labelled as ‘1 overlapping CI’.

Clustering of graphs was done by identifying the partition that maximized modularity. Relative degree scores for evading freezer and freezer networks were calculated using the following equation:

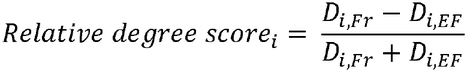

Where D_i,Fr_ and D_i,EF_ are the degree of the *i*th node for the freezer and evading freezer groups, respectively. Networks were visualized using Cytoscape version 3.10.2 (Shannon et al., 2003; Su et al., 2014).

### Coding and statistics

Analyses were performed using R version 4.3.3. To identify differences at specific time points during behavior we used FDR (Benjamini and Hochberg, 1995) corrected independent samples t-tests that compared CAS and vehicle treated groups. One sample t-tests were used to compare difference scores to zero with p-values FDR corrected. Missing values for brain activity were imputed via nuclear-norm regularized singular value decomposition as implemented in the softImpute package (version 1.4.3; Hastie et al., 2015; Mazumder et al., 2010). The regularization penalty (λ = 25) was determined empirically by identifying the inflection point in the scree plot of the singular values. Permutation resampling without replacement (10,000 times) was used to assess significance of strain and sex for each behavioral cluster. We used the igraph (Csardi and Nepusz, 2006; version 2.02) package for network analysis and the cccd (version 1.6) package to build the k-nearest neighbor networks.

## Funding

Funding was provided by the National Institute of General Medical Sciences (R35GM142566, JWK).

### Author contributions

Conceptualization: BDF and JWK; Methodology: BDF, JH, and JWK; Investigation: BDF, JH, NR, DK and DL; Formal analysis: BDF, NR and JWK; Writing: BDF and JWK.

### Data availability

All data needed to evaluate the conclusion in the paper are present in the paper and the supplemental material. Supplemental files are available at Dryad: https://doi.org/10.5061/dryad.sqv9s4nfh

## Acknowledgments

We thank Ian Lee for copy editing of the manuscript.

## List of supplementary material

*Supplemental figures*

Figure S2-1. Freezing behavior in response to different concentrations of CAS.

Figure S4-1. Internal clustering metrics for K-nearest neighbor partitions.

Figure S4-2. Behaviors associated with clustering using alternative variables.

Figure S4-3. Transition probabilities between behavioral states based on cluster.

Figure S6-1. Bootstrap ratios for PLS analysis.

Figure S6-2. Example covariations between cfos activity in individual brain regions and behavioral contrasts.

Figure S8-1. Functional network for non-reactive zebrafish.

Figure S8-2. Functional network for vehicle treated zebrafish.

*Supplemental tables*

Table S1-1. Parameters used for training random forest machine learning model.

Table S6-1. Brain region abbreviations.

*Supplemental files* (available at Dryad: 10.5061/dryad.sqv9s4nfh)

File S01. Statistics for Figure 2A.

File S02. Statistics for Figure 2B.

File S03. Behavioral data for Figure 2.

File S04. Neural *cfos* and behavioral data for Figure 6.

File S05. Bootstrap ratios for Figure 6.

File S06. Neural *cfos* data for network generation for Figure 7.

File S07. Correlations and confidence intervals for freezer, evading freezer, and non-reactive networks.

File S08. Graphml file for evading freezer network in Figure 8A.

File S09. Graphml file for freezer network in Figure 8A.

File S10. Graphml file for non-reactive network in Figure S8-1.

File S11. Graphml file for vehicle network in Figure S8-2.

File S12. Cytoscape file containing all four networks (evading freezer, freezer, non-reactive, and vehicle).

