## Supplemental figures for "Individual differences in fear memory expression engage distinct functional brain networks"

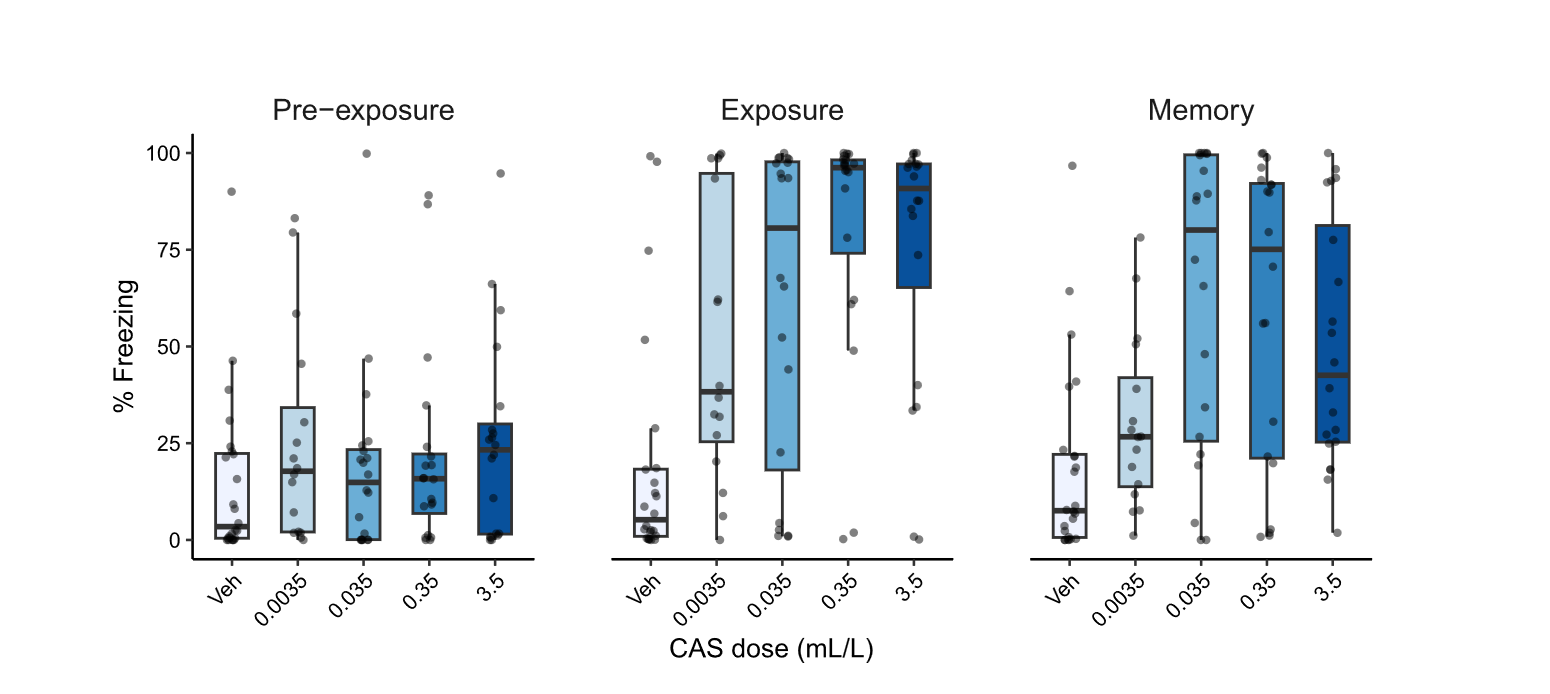
**Figure S2-1.** Freezing in response to different doses of CAS during pre-exposure, exposure, and memory day.

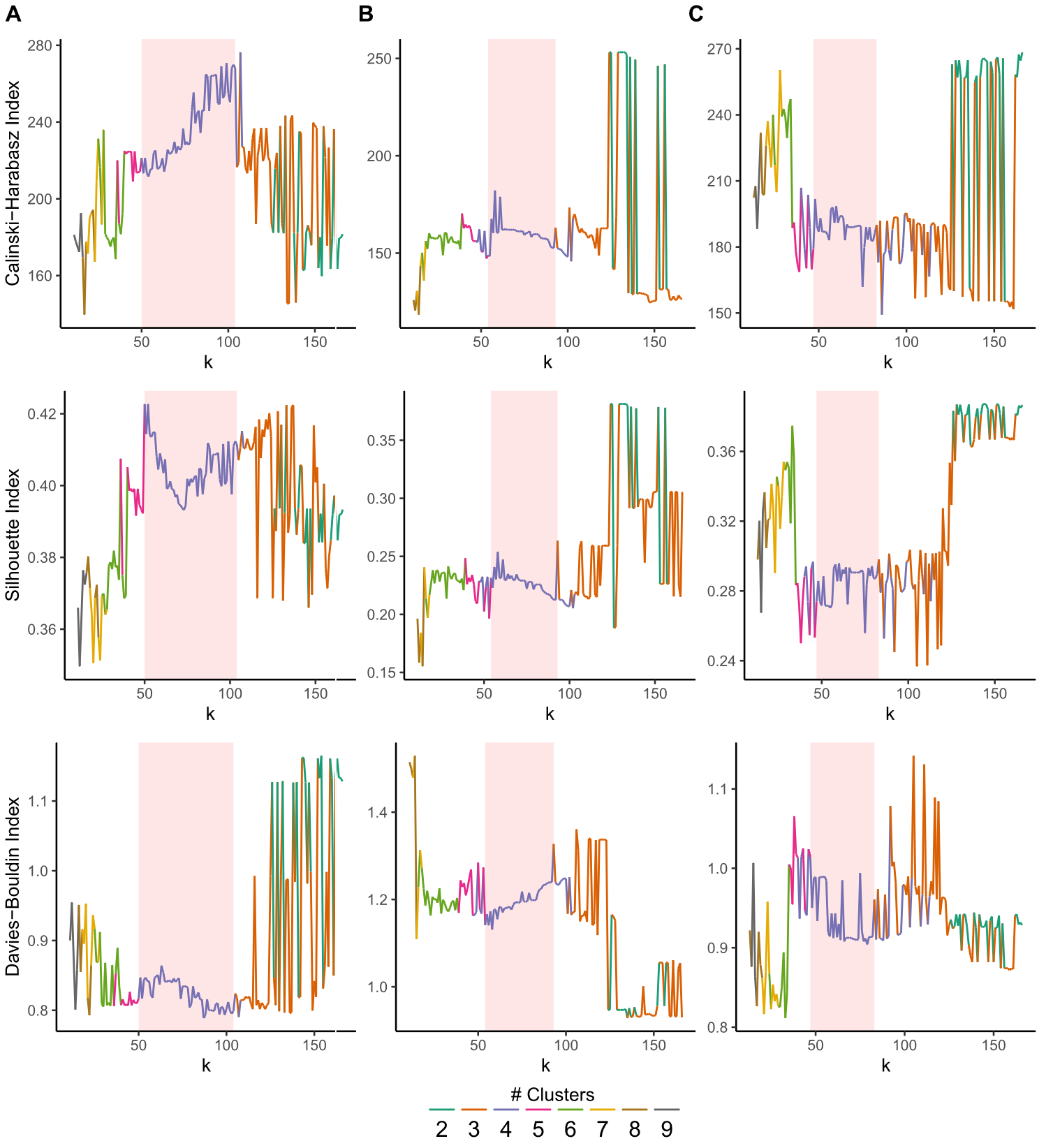

**Figure S4-1.** Internal clustering metrics applied to k-nearest-neighbor network partitions at different values of k for different variables used for clustering: (A) percent freezing, percent evasive, and percent evasive of active; (B) percent freezing and percent evasive; (C) isometric log ratios. Red shaded area marks regions highest stability over k. The midpoint of the shaded range was used for clustering.

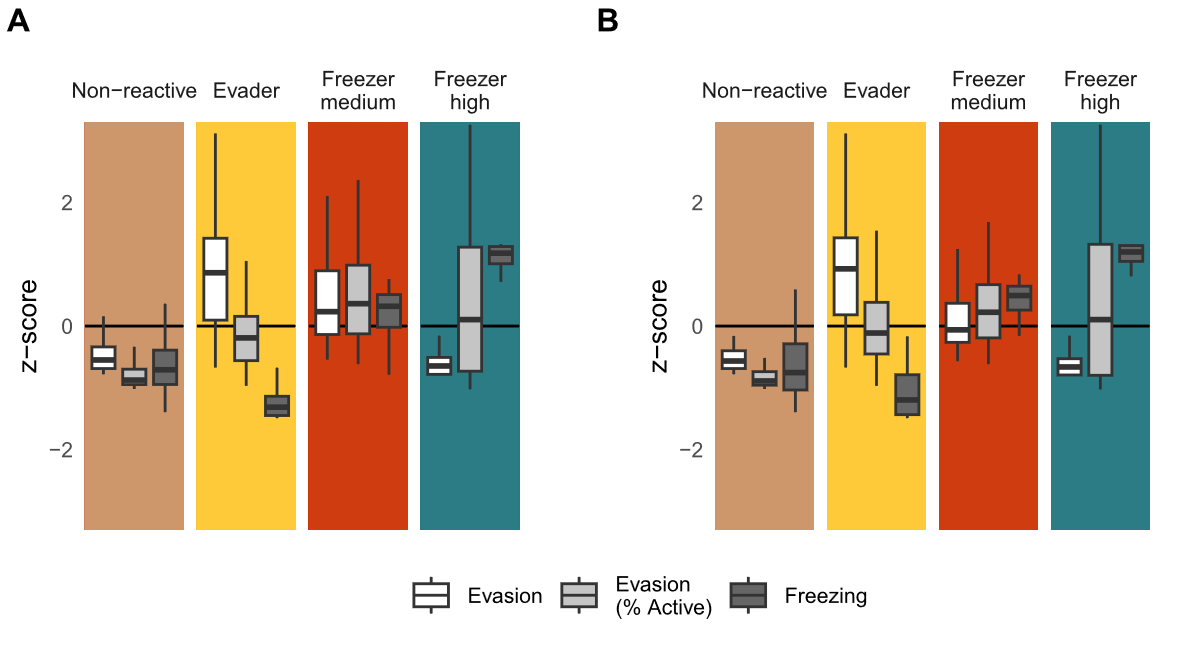

**Figure S4-2.** Behaviors associated with clustering using (A) percent freezing and percent evasion or (B) isometric log ratios.

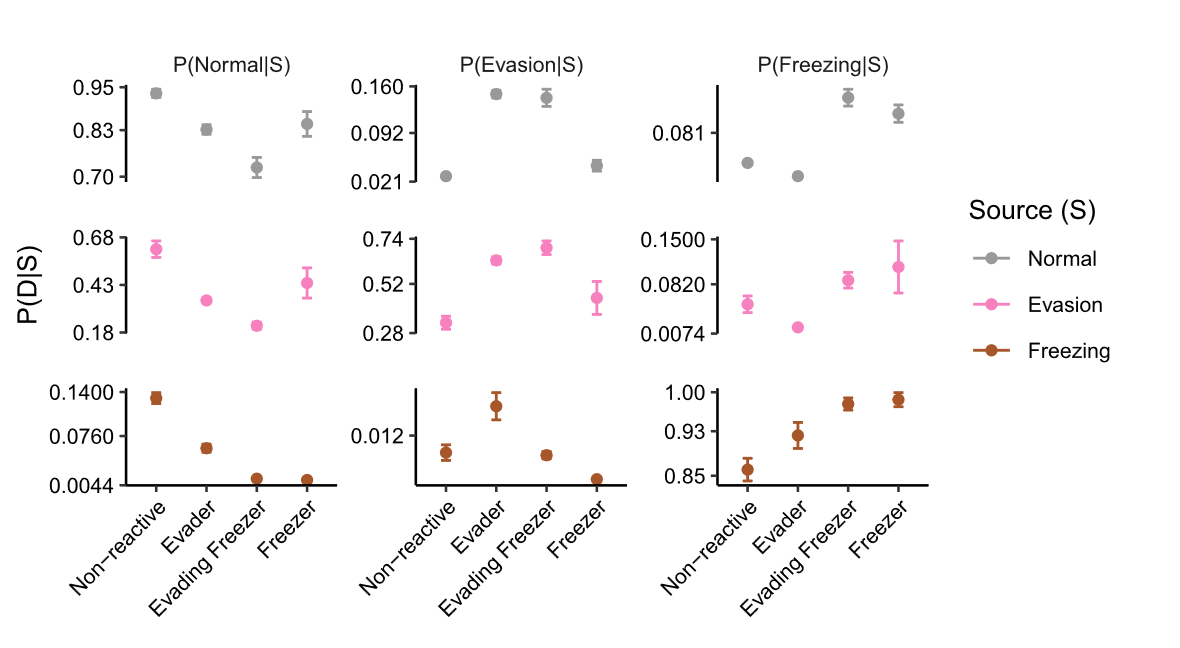

**Figure S4-3.** Transition probabilities between behavioral states based on behavioral cluster. D = destination, S = source. Error bars are 95% confidence intervals.

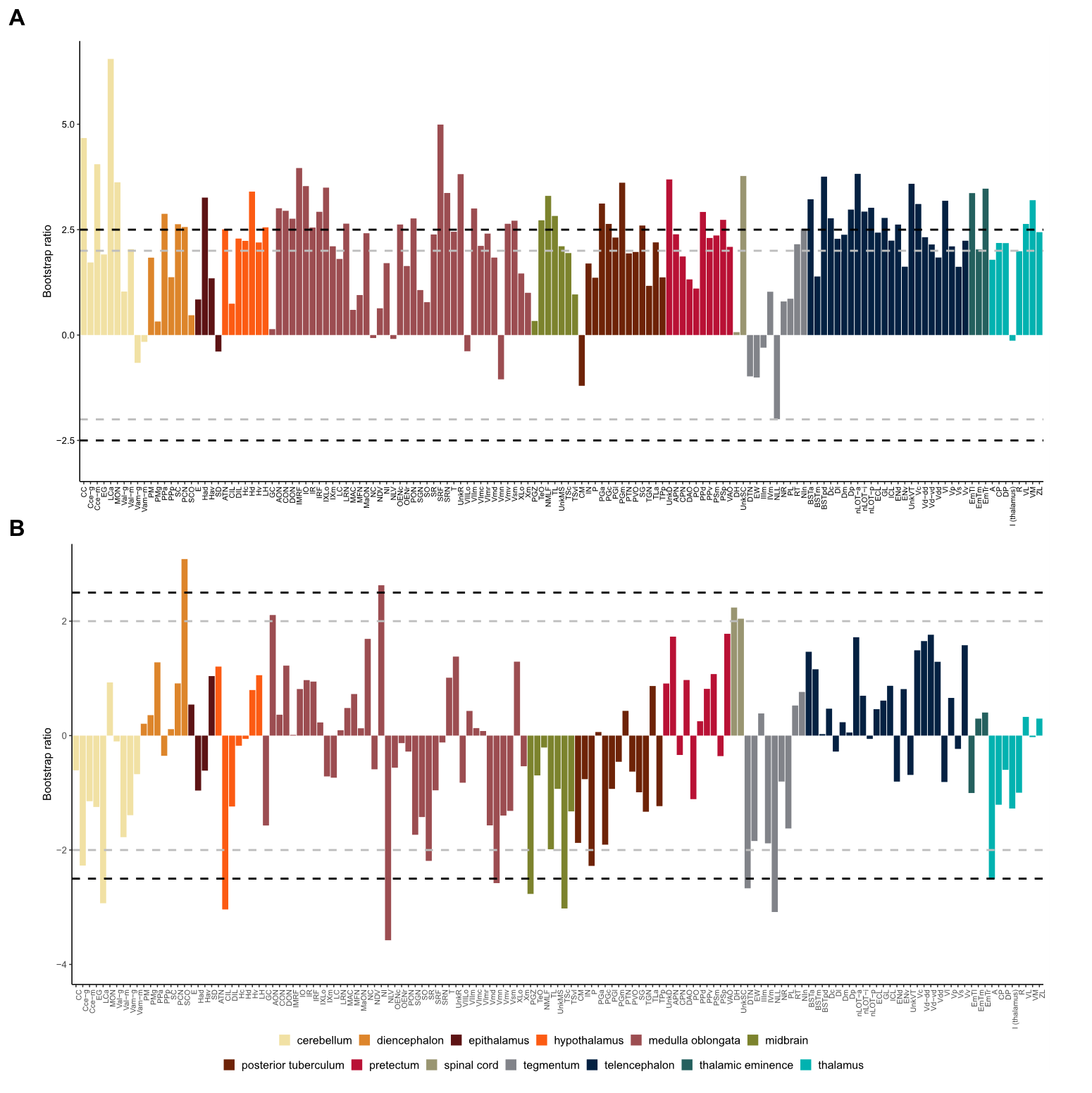

**Figure S6-1.** Bootstrap ratios from PLS analysis for the first (A) and second (B) contrasts. Region abbreviations can be found in Table S6-1.

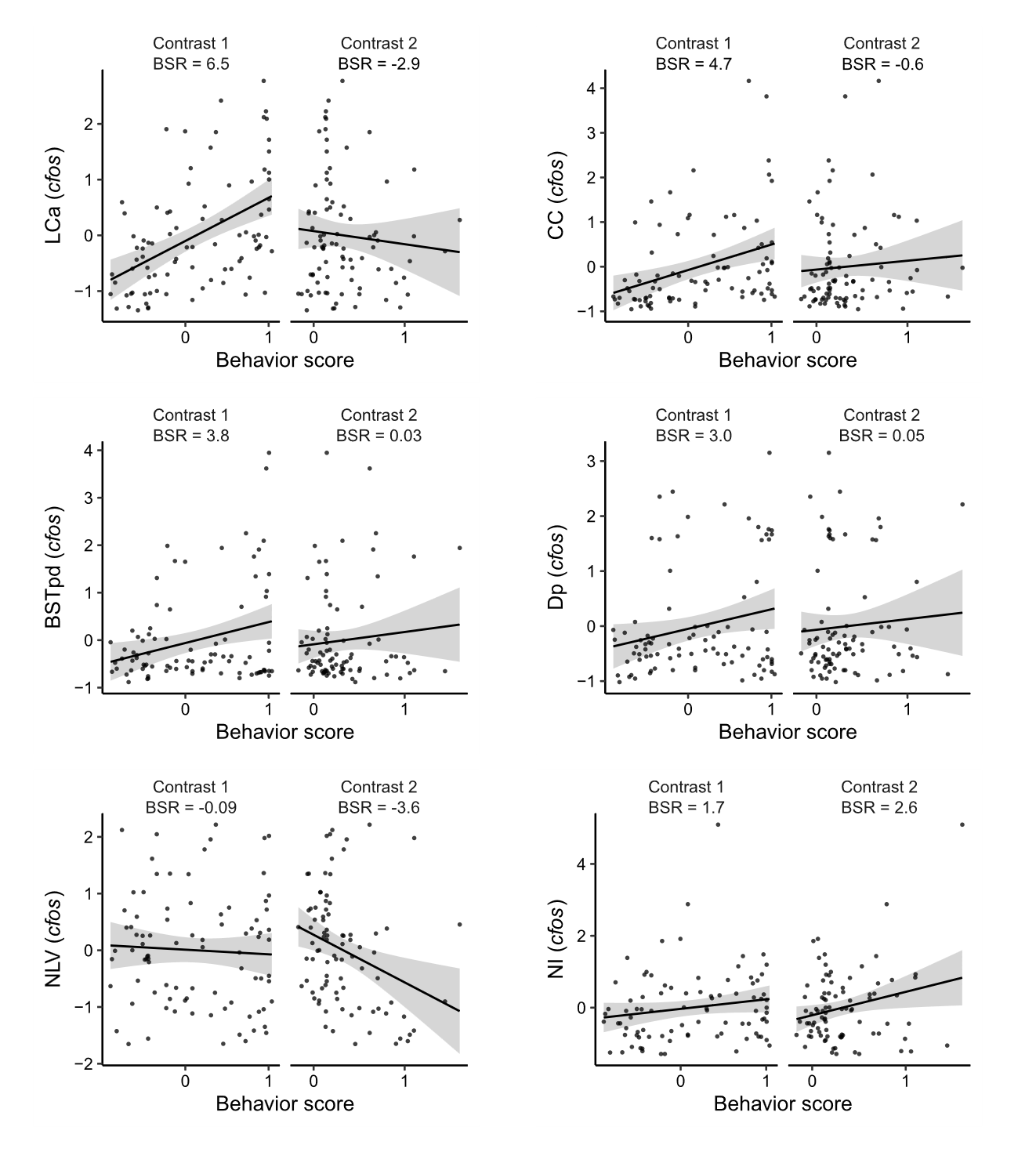

**Figure S6-2.** Example univariate relationships between standardized *cfos* activity behavioral scores related to contrasts 1 and 2 (Figure 6A). BSR = boot strap ratios calculated from the PLS analysis. Region abbreviations can be found in Table S6-1.

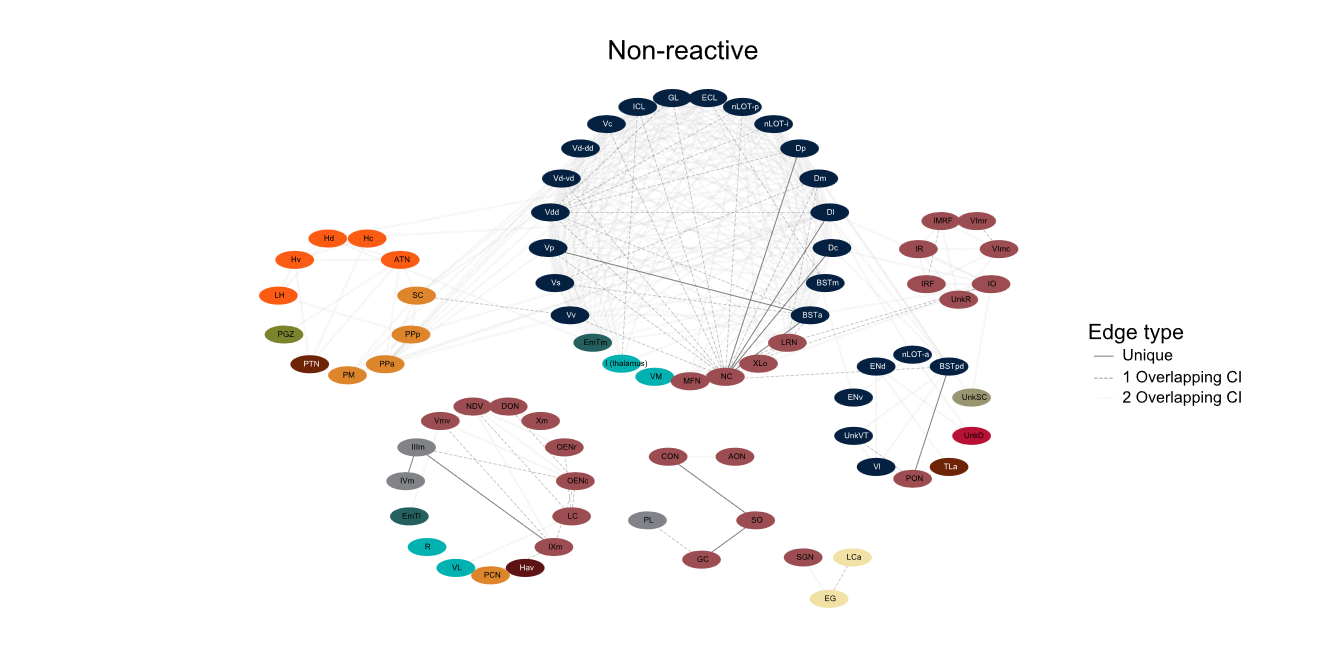

**Figure S8-1.** Functional network for non-reactive zebrafish. Nodes represent brain regions and edges represent suprathreshold correlations (FDR ≤ 0.001). Nodes follow the same coloring scheme as in Figure 7. Region abbreviations can be found in Table S6-1.

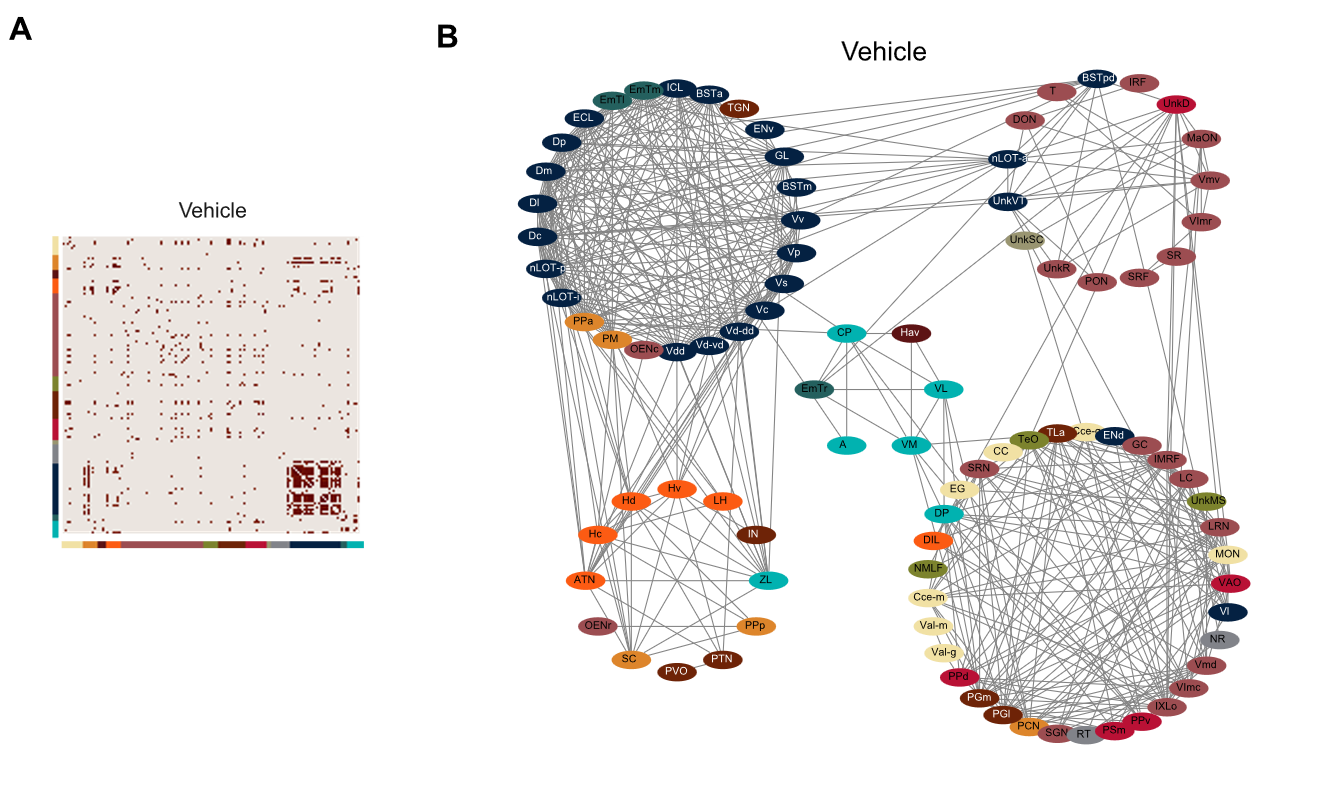

**Figure S8-2.** Functional network for vehicle treated zebrafish. A) Matrix of suprathreshold (FDR ≤ 0.001) correlations. B) Nodes represent brain regions and edges represent suprathreshold correlations. Nodes follow the same coloring scheme as in Figure 7. Region abbreviations can be found in Table S6-1.
